# One score to rule them all: regularized ensemble polygenic risk prediction with GWAS summary statistics

**DOI:** 10.1101/2024.11.27.625748

**Authors:** Zijie Zhao, Stephen Dorn, Yuchang Wu, Xiaoyu Yang, Alexander Beckerman, Jin Jin, Qiongshi Lu

## Abstract

Ensemble learning has become a cornerstone for improving the predictive accuracy of polygenic risk scores (PRS), and nearly all recent multi-ancestry PRS methods incorporate ensemble learning as a final step. However, existing ensemble approaches require individual-level genotype data for model training, which limits their real-world applications, especially in non-European populations without sufficient genomic samples. Here, we introduce a statistical framework for constructing regularized ensemble PRS that integrates a large number of candidate PRS models using only genome-wide association study summary statistics. Through extensive analyses across multiple traits and populations, we demonstrate that our method consistently outperforms state-of-the-art PRS approaches within and across ancestries. This framework presents “one score to rule them all” for its capability to enable seamless integration of newly developed PRS models with existing ones, providing a scalable and general solution for future PRS development and application.

## Introduction

With the continued success of genome-wide association studies (GWAS)^1,2^ and increasingly accessible summary statistics from these studies, genetic prediction efforts have generally focused on creating polygenic risk scores (PRS) that combine individually negligible but collectively substantial effects from millions of single nucleotide polymorphisms (SNPs) in GWAS summary data. Over the years, PRS methodology has evolved through improved model designs^3-9^, integration of functional genomic annotations^8,10-12^, and multi-ancestry/multi-trait joint modeling^12-14^. Due to the diverse genetic architecture of complex traits and moderate signal-to-noise ratio at current GWAS sample sizes, not all PRS models perform equally well, and no single method consistently outperforms others^15-17^. Ensemble learning is a strategy that trains a machine learning model to combine multiple learning algorithms for better predictive performance. In recent years, researchers have applied ensemble learning to integrate multiple PRS into an aggregated score with improved performance compared to any single PRS model^15,18-20^. Earlier work used linear regression or penalized regression to develop ensemble PRS^18,19^. More recently, super learning has been introduced as an omnibus approach for ensemble PRS construction^21-23^. It employs an “ensemble of ensemble” modeling strategy to achieve additional prediction gains from various PRS model designs and ensemble techniques^24,25^. In particular, these approaches have proven effective in multi-ancestry PRS applications^21-23,26-28^, leveraging the many PRS models optimized for each ancestry respectively to improve the predictive performance in the target population. Due to its apparent effectiveness, nearly every recent PRS method employs ensemble learning in some way.

Unsurprisingly, ensemble learning is data-demanding – ensemble model training requires a holdout dataset independent from GWAS samples. This creates a major hurdle for ensemble PRS application. Often, in practice, a summary-level GWAS dataset is all there is for PRS model training, particularly in non-European ancestries where individual-level data are scarce. Even as large biobank cohorts become increasingly accessible, GWAS of late-onset diseases or any outcomes associated with low study participation still rely on meta-analysis of numerous disease-ascertained cohorts^29^. Further, even when a dataset with individual-level genotype and phenotype information is available, it may have been included in the GWAS meta-analysis, making PRS ensemble learning impossible without overfitting. Therefore, researchers who wish to employ ensemble learning need to partition the valuable testing dataset, which is often small in size if it even exists, leading to insufficient PRS benchmarking and reduced statistical power in downstream applications^18,19,30^. This problem is further exacerbated in applications of super learning, where finer data partitioning is required to train a multi-level ensemble model. To avoid the need for individual-level holdout datasets, we recently introduced an approach (PUMAS) to fine-tune PRS^31^ and obtain a linear combination of multiple PRS models using only GWAS summary statistics^32^. Although it was an important proof of concept, the simple linear combination approach may yield problematic results when integrating a large number of PRS models. A summary statistics-based approach that can employ more advanced ensemble learning strategies to further improve PRS utility is thus naturally desired.

Here, we address these challenges by introducing two summary statistics-based ensemble learning techniques, which we have incorporated into the PUMAS-ensemble software suite^32^. Our elastic net (PUMAS-EN) ensemble learning approach simultaneously conducts ensemble model training, fine-tuning, and benchmarking based on a large number of input PRS models. We also introduce a super learning approach (PUMAS-SL) to combine multiple regularized ensemble PRS. Through extensive simulations and analysis of many datasets, we demonstrate the robust and superior performance of our approach both within and across ancestries. Most importantly, we showcase the capability of our approach to continuously integrate new PRS models with many existing ones, which makes it a general framework that can self-evolve by always including newly-introduced PRS models from the literature.

## Results

### Method overview

We first provide an overview of our workflow and will delve into statistical details in the **Methods** section. If a dataset with individual-level genotype and phenotype information is available, the conventional strategy for fitting and evaluating ensemble PRS models is to split the full sample into independent subsets for ensemble model training and benchmarking (**Fig. 1A**). If the ensemble model has tuning parameters, e.g., regularization parameters in penalized regressions, the dataset for model training needs to be further divided so that a subset can be used for hyperparameter fine-tuning. However, since an individual-level dataset with sufficient samples is often unavailable in practice, we present an ensemble learning process following a similar modeling framework but requiring only GWAS summary statistics. In PUMAS-ensemble, we partition the full GWAS summary statistics dataset into three down-sampled summary statistics datasets for training, ensemble learning, and testing, respectively (**Fig. 1B**). With these down-sampled summary statistics datasets, we can train multiple PRS models by various methods, apply the two ensemble learning approaches based on elastic net and super learning to integrate these PRS models, and finally, benchmark the performance of the constructed ensemble PRS. Only GWAS summary statistics and linkage disequilibrium (LD) reference data are required in this framework. We note that this is a general framework that allows researchers to choose and combine any set of PRS methods for improved prediction. For illustration, we considered eight common PRS methods for most of our analyses in this study^4-9,33-35^ (**Supplementary Table 1**). All details of method implementation are presented in **Methods**.

**Fig. 1.**
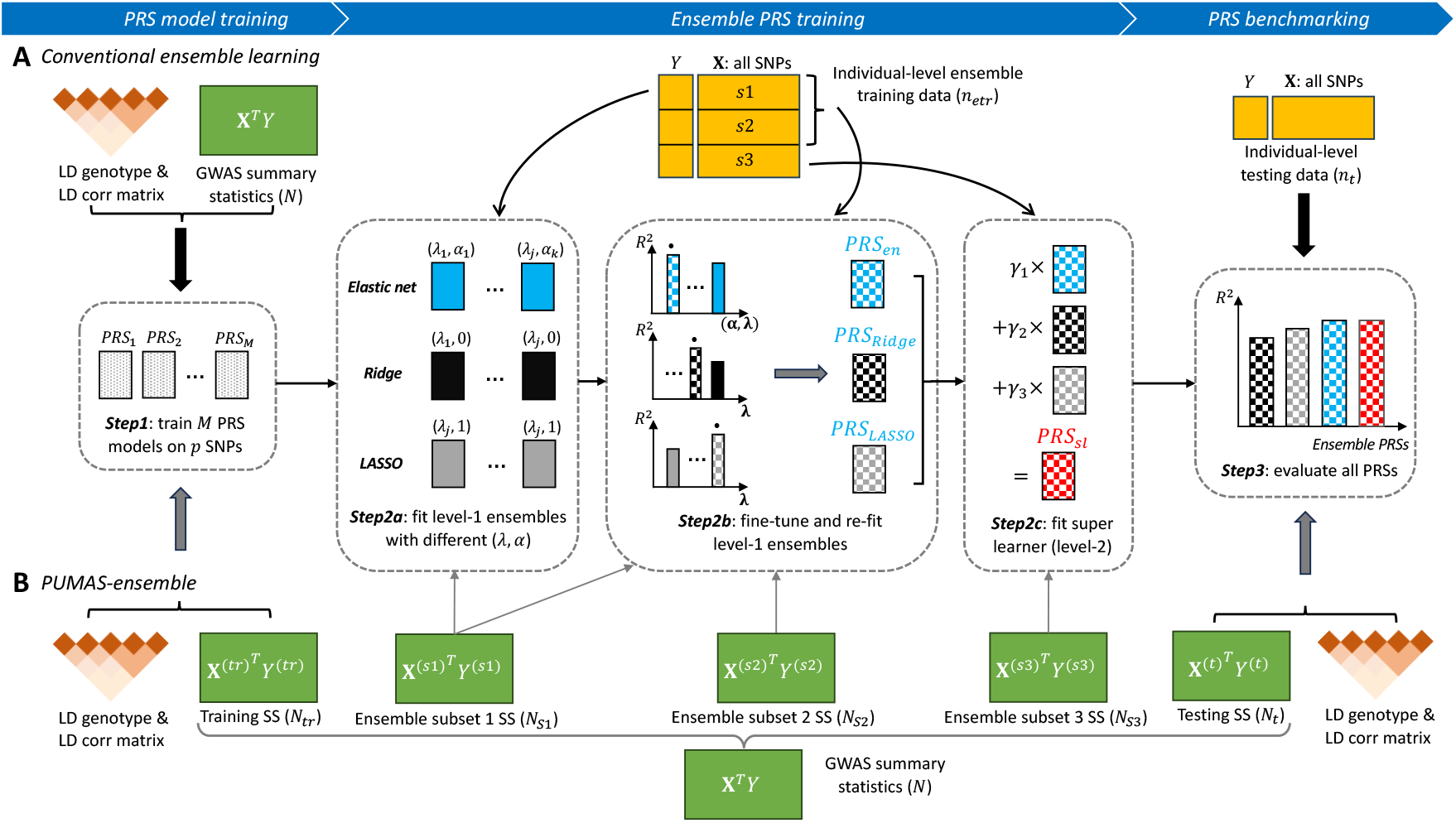
Workflow of summary-statistics-based ensemble learning. (**A**) Conventional ensemble learning approaches require individual-level genotype and phenotype data (orange boxes) to train ensemble learning models and evaluate predictive performance. (**B**) The proposed PUMAS-ensemble approach can follow the same procedure without the need for individual-level data. It leverages a resampling strategy to partition the full GWAS summary statistics into multiple sub-sampled summary datasets for different analytical aims.

### Simulation studies

We randomly selected 100,000 independent individuals of European descent in UK Biobank (UKB)^36^ and utilized their genotype data on 944,547 HapMap 3 SNPs in our simulation study. To emulate diverse genetic architectures, continuous trait values were generated based on different levels of heritability and different numbers of causal variants (**Methods**). We compared the performance of three ensemble strategies: linear regression, elastic net, and super learning, using either the conventional individual-level data-based ensemble learning and summary-level data-based PUMAS-ensemble. For conventional ensemble learning, we partitioned the full individual-level dataset into three subsets for single PRS training (*N*_*tr*_=60,000), ensemble training (*N*_*en*_=30,000), and testing (*N*_*t*_=10,000), respectively. Similarly, for PUMAS-ensemble, we performed GWAS to obtain summary statistics based on the full sample, and then subsampled three sets of summary statistics to train single PRS models (*N*_*tr*_=60,000), combine them to obtain the ensemble PRS (*N*_*en*_=30,000), and evaluate performance (*N*_*t*_=10,000). A total of 110 single PRS models were combined by each ensemble PRS method. For both individual-level and summary-level analyses, we performed 4-fold Monte Carlo cross-validation (MCCV)^31,32^ and reported the average predictive *R*^2^.

Both the elastic net-based and super learning-based ensemble PRS approaches (i.e., PUMAS-EN and PUMAS-SL) showed superior predictive performance compared to the single PRS models under all simulation settings (**Fig. 2; Supplementary Table 2**). Using the median accuracy of single PRS models as the baseline, PUMAS-EN and PUMAS-SL improved predictive *R*^2^ by 11.9%-22.0% and 10.7%-23.2%, respectively, across the various simulation settings, demonstrating robust and nearly identical performance. Notably, summary-statistics-based elastic net and super learning showed highly consistent results compared to the conventional ensemble learning based on individual-level data. Linear regression ensemble learning without regularization was affected by substantial collinearity among PRS models and showed poor results, especially under externally estimated LD in summary-statistics-based inference. Regularized ensemble models maintained stable performance as additional correlated models were added to the ensemble inputs, whereas the linear regression ensemble degraded further (**Supplementary Figure 1**). These results highlight the importance of statistical regularization in summary-statistics-based ensemble learning, especially when aggregating a large number of PRS models. Because of this observation, we will focus on elastic net and super learning in following analyses.

**Fig. 2.**
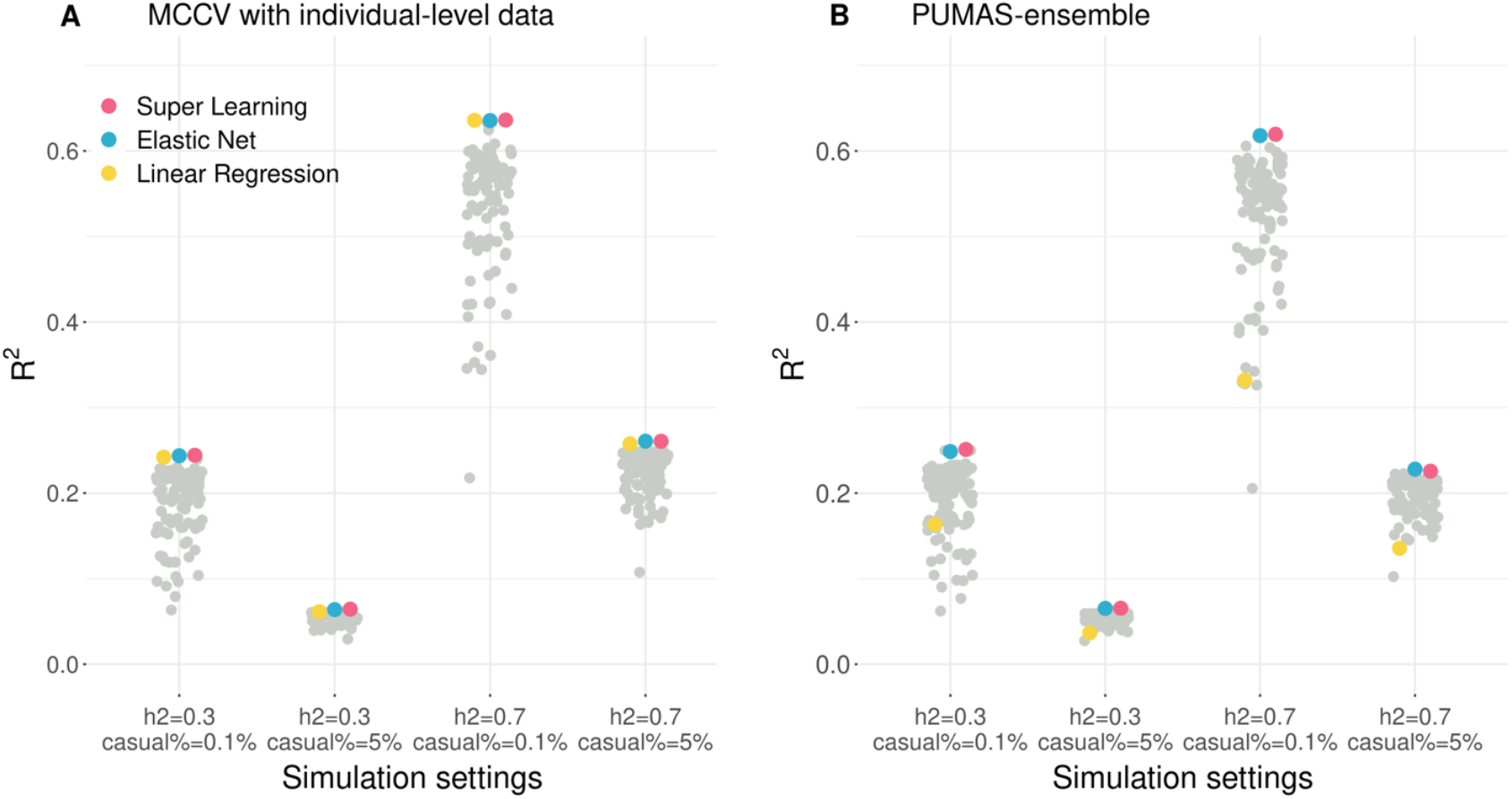
Performance of various ensemble PRS strategies on simulated data. (**A**) Performance of ensemble PRS trained on individual-level data. Prediction accuracy was estimated based on 4-fold MCCV. (**B**) Performance of PUMAS-ensemble PRS quantified by summary-statistics-based 4-fold MCCV. Ensemble PRS models, i.e., linear regression, elastic net, and super learning, are highlighted. Single PRS models are shown in gray. X-axis: simulation settings; Y-axis: predictive *R*^2^; h2: heritability; causal%: proportion of causal variants. Detailed simulation results are summarized in **Supplementary Table 2**.

### Ensemble PRS outperforms single PRS models in UKB

Next, we applied PUMAS-ensemble to 16 complex traits in UKB (**Supplementary Table 3**) and compared its prediction accuracy to the conventional ensemble learning strategy based on individual-level data. Specifically, we trained conventional ensemble PRS on three quarters of a holdout UKB dataset (N=38,521) and benchmarked all PRS models on the remaining quarter of samples (**Methods**). Similar to what we observed in simulations, both PUMAS-EN and PUMAS-SL showed consistently higher predictive R^2^ compared to single PRS models trained on the same dataset (**Fig. 3A; Supplementary Table 4**). Summary-statistics-based elastic net and super learning respectively achieved 31.3% and 30.6% average increase in predictive *R*^2^ across 16 traits compared to the median performance of the 110 single PRS models. Compared to the tuning-free PRS models such as LDpred2-auto^6^ and PRS-CS-auto^5^, PUMAS-EN had a 9.7% and 20.6% average increase in R^2^, respectively, across the 16 traits. Similarly, PUMAS-SL had a 9.2% and 20.0% average increase in R^2^, respectively. This suggests that the substantial performance gain of PUMAS-ensemble is achieved by ensemble learning of multiple PRS models rather than improved fine-tuning of single PRS models.

**Fig. 3.**
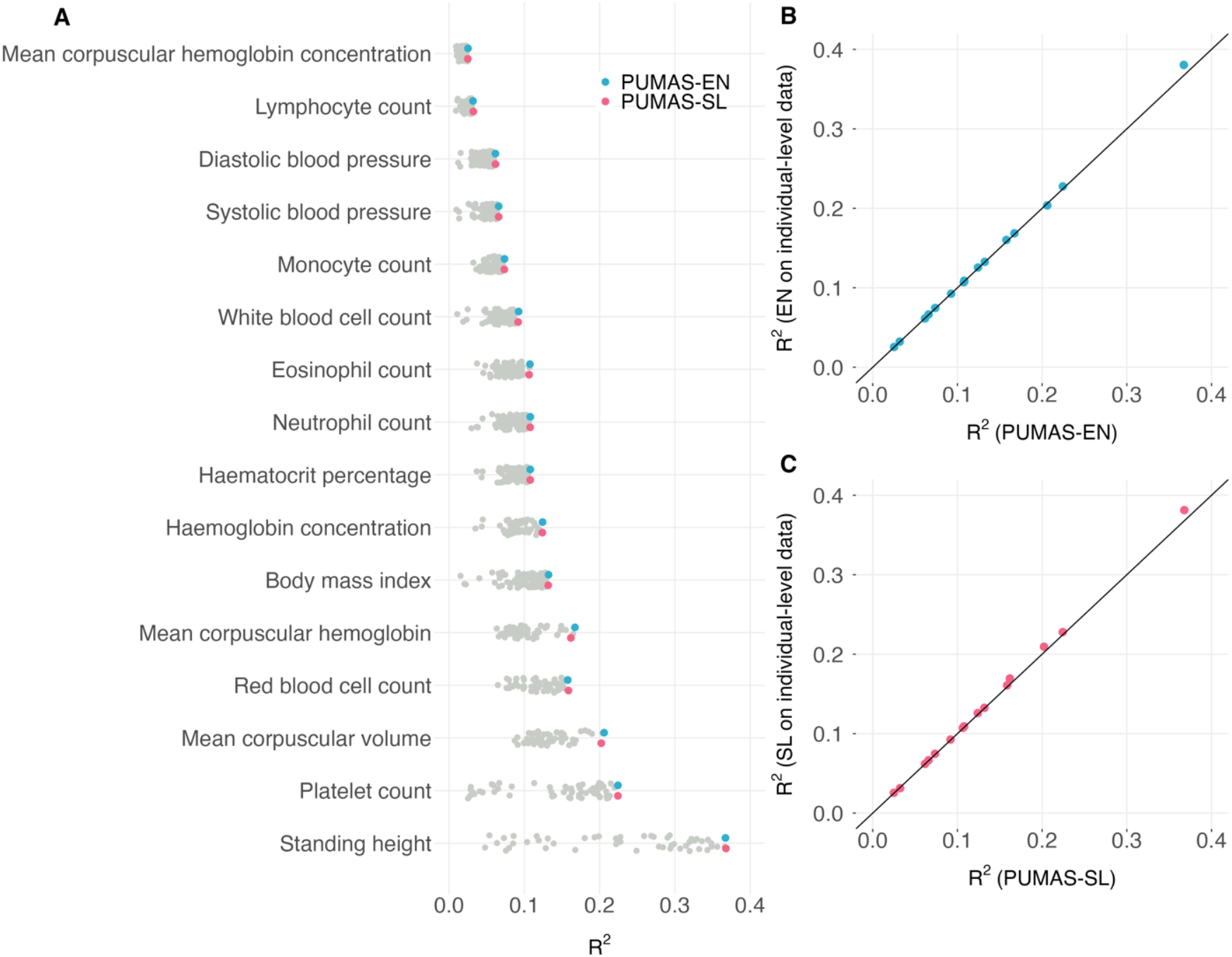
Ensemble PRS prediction for 16 traits in UKB. (**A**) Comparison of PUMAS-ensemble PRS with single PRS models on the holdout UKB dataset (N=38,521). Single PRS models are colored in gray while PUMAS-EN and PUMAS-SL scores are highlighted in blue and red, respectively. Y-axis: trait names; X-axis: predictive *R*^2^. (**B**-**C**) Comparison of summary-statistics-based ensemble PRS with elastic net (panel B) and super learning (panel C) PRS trained on holdout UKB data. Y-axis: *R*^2^ of conventional ensemble PRS; X-axis: *R*^2^ of PUMAS-ensemble PRS. The diagonal line indicates equal *R*^*2*^ by PUMAS-ensemble PRS and conventional ensemble PRS. Details of the UKB GWAS summary statistics used for PRS training are summarized in **Supplementary Table 3**. Predictive performance of all models and traits is reported in **Supplementary Table 4**.

In addition, the performance of PUMAS-ensemble PRS was almost indistinguishable from the ensemble PRS trained based on individual-level data (**Fig. 3B-C; Supplementary Table 4**). We also benchmarked against PRSmix^37^ which implements an elastic net ensembling strategy with additional filtering of input PRS, and obtained similar results (**Supplementary Figure 2A-B**). In practice, individual-level holdout datasets often have smaller sample sizes when they do exist. When we reduced the size of individual-level ensemble training data to N=500, PUMAS-EN and PUMAS-SL outperformed elastic net and super learning by an average R^2^ increase of 15.6% and 22.8%, respectively (**Methods; Supplementary Table 4**). These findings suggest that GWAS summary-level data alone is sufficient for building powerful ensemble PRS in real-world applications, and can even outperform conventional ensemble learning when the individual-level holdout dataset has limited sample size.

When an individual-level holdout dataset does not exist, unsupervised learning methods such as PRS-PCA^38^ may provide an alternative strategy to integrate multiple PRS models. We benchmarked PUMAS-ensemble’s performance to PRS-PCA (**Methods; Supplementary Table 4**), and found significantly better performance of PUMAS-EN (6.60%; p-value=1.5e-05; Wilcoxon signed-rank test) and PUMAS-SL (6.11%; p-value=1.5e-05), highlighting the need to employ supervised learning for optimal predictive performance.

As for computation, using height data as an example (51 PRS models for 1.2M SNPs), PUMAS-EN and PUMAS-SL took 1.85 minutes and 2.57 minutes, respectively. Additionally, PUMAS-ensemble’s runtime (4 CPU threads) increased linearly as the number of input PRS increased, while its memory usage scaled at a sublinear rate (**Methods; Supplementary Table 5**). These results demonstrate that PUMAS-ensemble is a computationally scalable framework for combining hundreds of scores in routine PRS analysis.

### PUMAS-ensemble PRS demonstrates robust out-of-sample performance in AllofUs

Additionally, we investigated the out-of-sample performance of ensemble PRS. We applied PUMAS-ensemble to build PRS for standing height, body mass index, and six binary traits including bipolar disorder, major depressive disorder, schizophrenia, breast cancer, coronary artery disease, and type II diabetes using well-powered and publicly available GWAS summary-level datasets^39-46^. Details on these GWAS datasets are presented in **Supplementary Table 6**. We extracted corresponding phenotypes and evaluated PRS prediction accuracy in the All of Us Research Program^47^ (AllofUs) (**Methods; Supplementary Table 7**). We compared PUMAS-EN, PUMAS-SL, and single PRS models on independent AllofUs participants of European ancestry and reported prediction *R*^2^ for continuous traits and AUC for binary traits. We observed consistent performance between PUMAS-EN and PUMAS-SL (**Fig. 4; Supplementary Tables 8-9**). On average, summary-level elastic net and super learning ensemble PRS improved prediction accuracy by 16.75% and 16.79% for continuous traits compared to the median *R*^2^ of the 110 single PRS models, and 6.49% and 6.37% for disease outcomes compared to the baseline-adjusted median AUC (**Methods**). PUMAS-ensemble had nearly identical performance to individual-level data-based ensemble learning, demonstrating consistency for clinical outcomes (**Supplementary Figure 2C-D)**.

**Fig. 4.**
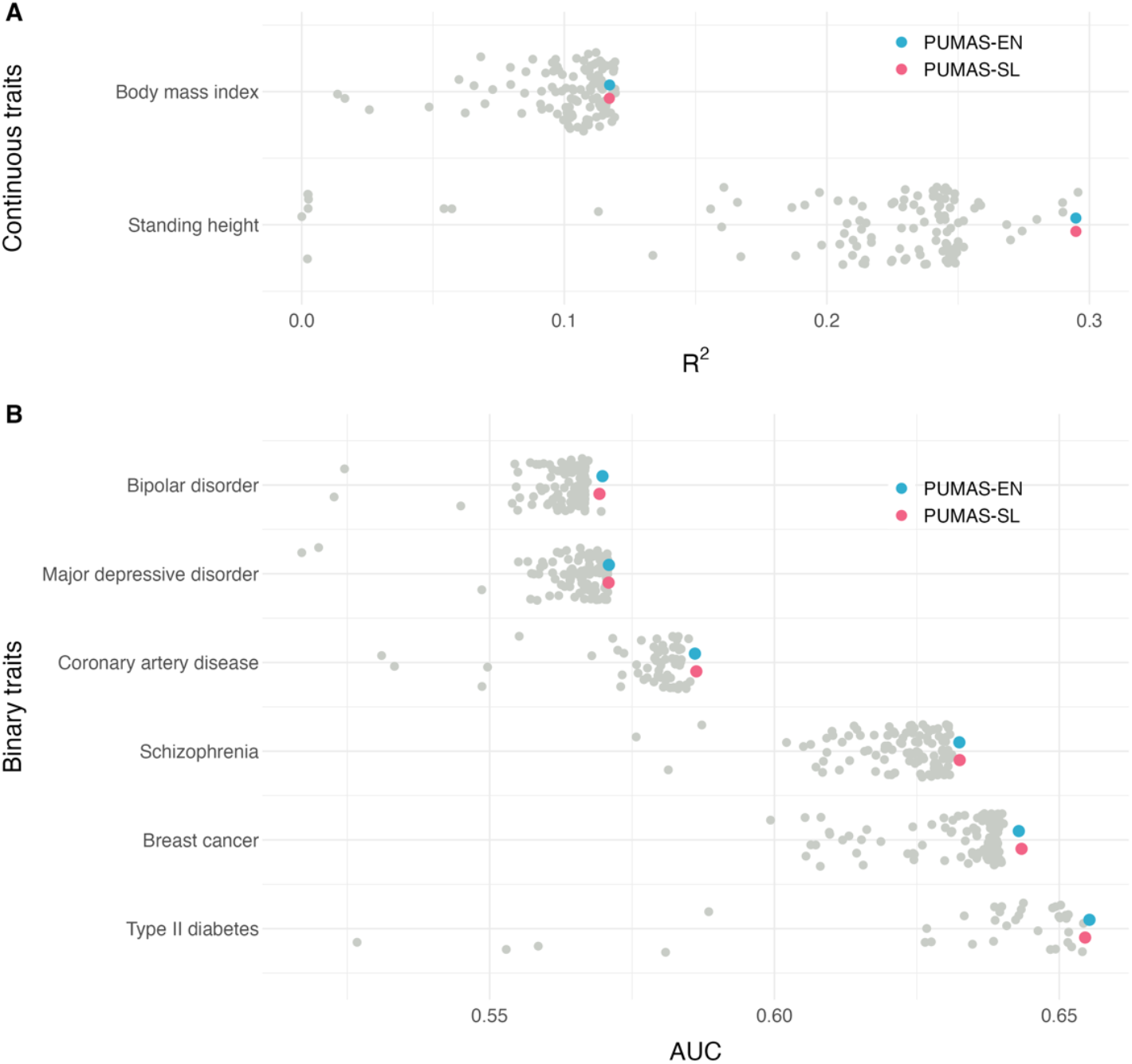
Evaluation of PUMAS-ensemble PRS in AllofUs. Predictive performance of PUMAS-EN, PUMAS-SL, and single PRS models are shown in blue, red, and gray, respectively. PRS Performance is shown for both **(A)** continuous traits with predictive *R*^2^ and **(B)** binary traits with AUC (without covariates). X-axis: *R*^2^ or AUC; Y-axis: phenotype. The list of included phenotypes and detailed results are summarized in **Supplementary Tables 6**-**9**. AUC: area under the receiver operating characteristic curve.

### Cross-ancestry ensemble PRS improves polygenic prediction accuracy on East Asian individuals in AllofUs

Next, we showcased the benefit of employing our ensemble PRS strategy in a cross-ancestral risk prediction setting. We extended PUMAS-ensemble to construct optimal ensemble scores for participants of East Asian (EAS) descent in AllofUs by aggregating PRS models trained using European (EUR) GWAS summary data from UKB and EAS GWAS summary data from Biobank Japan (BBJ)^48-50^ (**Methods; Fig. 5A**). We then compared PUMAS-EN and PUMAS-SL with the single EUR and EAS PRS models on four traits that are well-powered in both UKB GWAS and BBJ GWAS: standing height, body mass index, diastolic blood pressure, and systolic blood pressure (**Supplementary Table 10**). The ensemble approach generates the best-performing PRS model for all traits analyzed, with markedly improved prediction accuracy comparing to single-ancestry PRS models (**Fig. 5B-E; Supplementary Table 11**). Across the four traits, predictive *R*^2^ of PUMAS-EN is 70.7% and 179.5% higher on average than the median *R*^2^ of the various EAS and EUR PRS models (68.6% and 179.1% for PUMAS-SL), respectively. Additionally, we compared PUMAS-ensemble performance to cross ancestry PRS method PRS-CSx run without an additional validation cohort. Predictive *R*^2^ of PUMAS-EN is 55.0% higher on average than the predictive *R*^2^ from cross-ancestry method PRS-CSx (52.5% for PUMAS-SL). Between the two proposed ensemble strategies, PUMAS-EN and PUMAS-SL have quite consistent performance with similar average R^2^ across the four traits. PUMAS-ensemble continued to have identical performance to individual-level data-based ensemble learning in this cross-ancestry application (**Supplementary Figure 2E**).

**Fig. 5.**
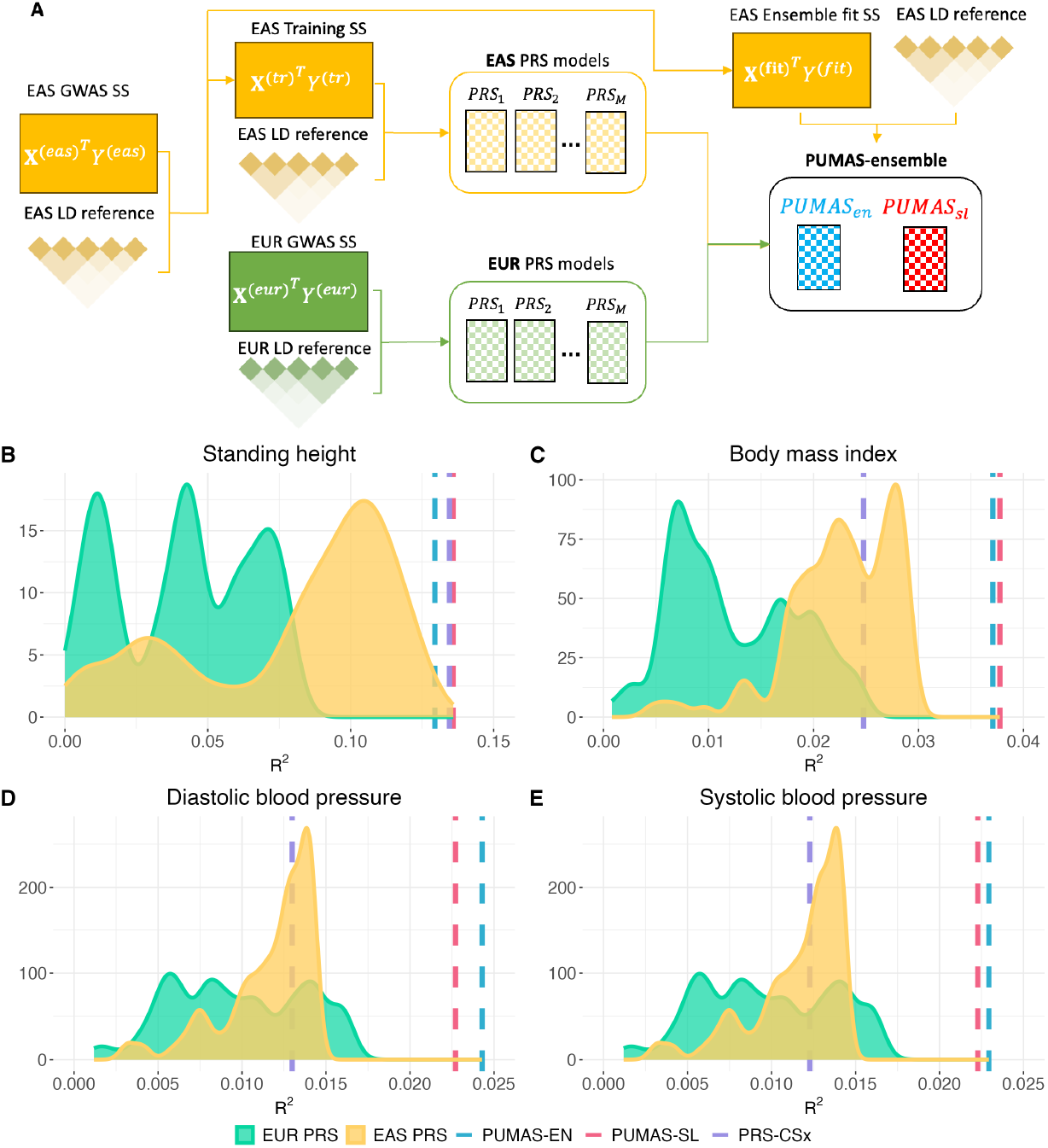
Cross-ancestral performance of PUMAS-ensemble PRS in AllofUs. **(A)** Workflow for cross-ancestry PUMAS-ensemble application. **(B-E)** We trained PUMAS-EN and PUMAS-SL PRS models for **(B)** standing height, **(C)** body mass index, **(D)** diastolic blood pressure, and **(E)** systolic blood pressure, combining single EUR PRS and EAS PRS, i.e., PRS models trained based on either EUR GWAS from UKB or EAS GWAS from BBJ. Model performance was evaluated on EAS participants in AllofUs. The R^2^ distribution of single PRS models trained based on EUR and EAS GWAS are shown in yellow and green, respectively. R^2^ of PUMAS-EN and PUMAS-SL are highlighted as blue and red dashed lines. R^2^ of PRS-CSx is highlighted as a purple dashed line. Y-axis: number of PRS models; X-axis: predictive *R*^2^. Full results for all models and traits are reported in **Supplementary Table 11**.

### Cross-ancestry ensemble PRS of blood lipid traits for multiple ancestries

We further evaluated the performance of our ensemble PRS method on blood lipid traits across four ancestries on validation individuals from UKB. We utilized ancestry-stratified GWAS summary data from the Global Lipids Genetics Consortium^51^ (GLGC) for high-density lipoprotein (HDL) cholesterol, low-density lipoprotein (LDL) cholesterol, log-transformed triglycerides (logTG), and total cholesterol (TC), across four ancestries, including African (AFR), EAS, EUR, and South Asian (SAS) populations (**Supplementary Table 12**). For each genetic ancestry group, we used PUMAS-ensemble to generate scores for individuals of a matching ancestry in UKB by aggregating models trained using subsampled GLGC summary data from a matching ancestry and full GLGC summary data from the other three ancestries (**Supplementary Figure 3**). The consortium also provides data for Admixed American and Hispanic/Latino population (AMR); however, we did not include them in our analysis because two traits were flagged during our GWAS QC process and there is limited sample size of AMR ancestry individuals in UKB (**Methods**). We compared PUMAS-EN and PUMAS-SL with PRS methods of single ancestry (**Fig. 6; Supplementary Table 13**). For all traits, the ensemble approach outperforms the median *R*^2^ of single PRS models of the same ancestry by an average of 370.0% for PUMAS-EN (median of 298.7%) and 370.7% for PUMAS-SL (median of 299.4%). Compared to the median *R*^2^ of single PRS models of European ancestry, PUMAS-EN outperforms by an average of 132.0% (median of 101.1%) and PUMAS-SL by 133.0% (median of 101.1%). PUMAS-ensemble outperforms PRS-CSx^27^ by an average of 148.9% for PUMAS-EN (median of 31.4%) and 148.5% for PUMAS-SL (median of 33.5%) (**Supplementary Table 14; Supplementary Figure 4**). We note that PUMAS-EN and PUMAS-SL again delivered comparable prediction accuracy compared to individual-level data-based ensemble learning (i.e, PRSmix^37^), exemplified by Pearson correlations of 0.979 (p-value=4.57e-11) and 0.983 (p-value=1.04e-11), respectively (**Supplementary Table 14; Supplementary Figure 2F-G**). This highlights the ability of our summary data-based ensemble PRS approach in cross-ancestry applications to improve over single PRS models and to leverage the large sample size of European GWAS to enhance prediction accuracy in diverse non-European populations.

**Fig. 6.**
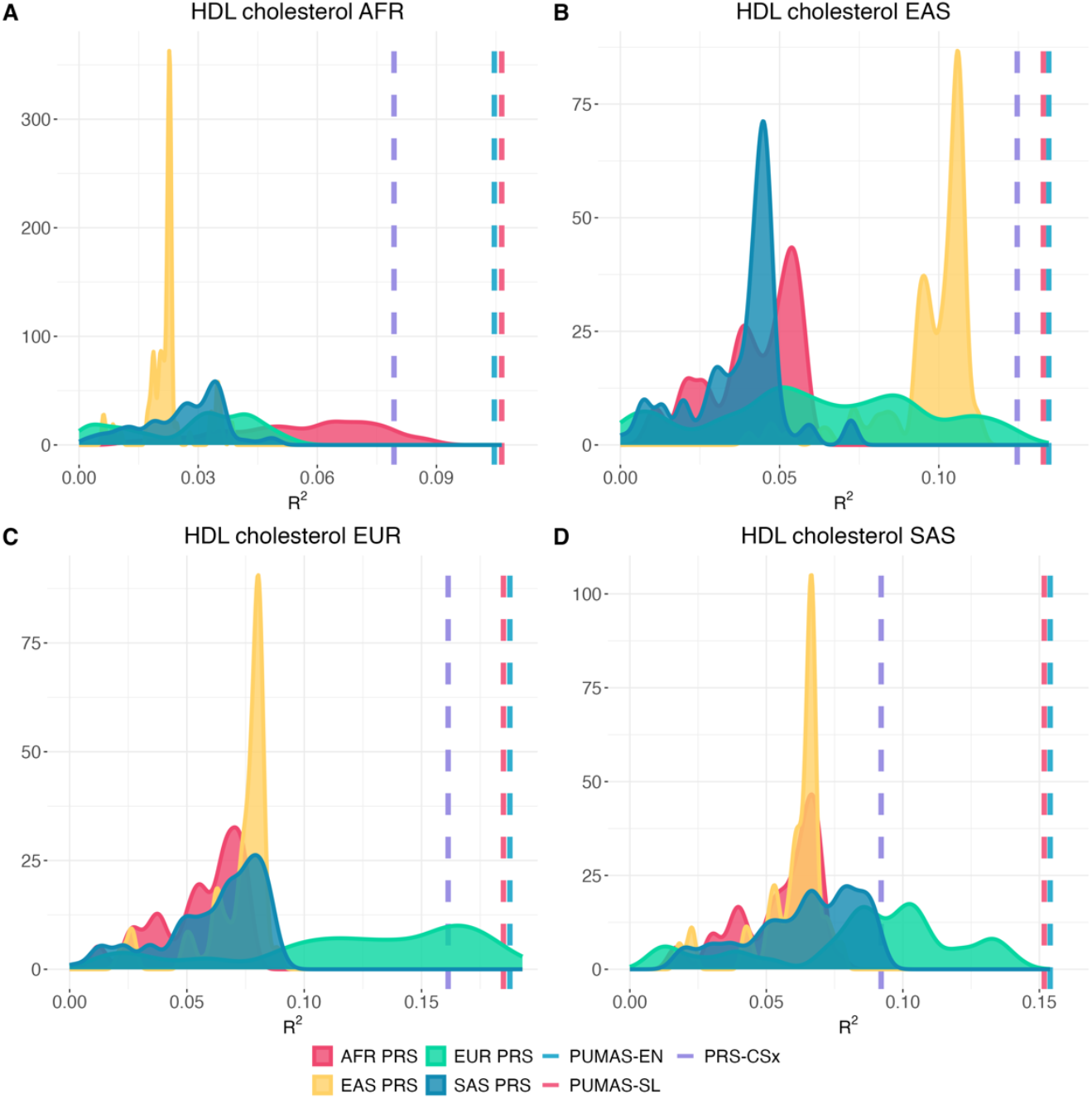
Cross-ancestry performance of PUMAS-ensemble PRS on HDL cholesterol. **(A-D)** We trained PUMAS-EN and PUMAS-SL PRS models for HDL cholesterol for ancestries **(A)** AFR, **(B)** EAS, **(C)** EUR, and **(D)** SAS, combining single PRS models trained based on ancestry-specific GWAS from GLGC. Model performance was evaluated on the highlighted ancestries participants in UKB. The R^2^ distribution of single PRS models trained based on AFR, EAS, EUR, and SAS ancestries are highlighted in pink, yellow, green, and blue, respectively. The R^2^ of PUMAS-EN, PUMAS-SL, and PRS-CSx are marked by dashed lines. Y-axis: number of PRS models; X-axis: predictive *R*^2^. Full results for all models and traits are reported in **Supplementary Table 13**.

It is well established that LD difference between populations can be a limiting factor for PRS transferability across genetic ancestries^52,53^. Here, we further investigate the impact of LD misspecification on PUMAS-ensemble for cross-ancestry prediction. We re-trained and evaluated PUMAS-ensemble using 1000 Genomes Project LD data (ancestry-specific panels) instead of the UKB LD reference in the ensemble training step (**Methods**). We found that using target-sample LD (e.g., UKB EUR LD for EUR samples in UKB) showed overall consistent PRS accuracy compared to using ancestry-matched external LD (e.g., 1KG EUR LD for EUR samples in UKB) for both PUMAS-EN (0.77% average improvement) and PUMAS-SL (0.2% average improvement) (**Supplementary Table 15; Supplementary Figure 5**). This suggests that it is acceptable to use external LD datasets with matched genetic ancestry in ensemble PRS applications. This observation is consistent with our earlier work^32^ as well as several other recent papers^30,54^. On the other hand, using population-mismatched LD (e.g., 1KG EUR LD for EAS samples in UKB) resulted in average reduction of ensemble performance by 13.0% and 14.0% for PUMAS-EN and PUMAS-SL, respectively (**Supplementary Table 15; Supplementary Figure 6**). These results reaffirm that ancestral concordance between the LD, GWAS, and target datasets is critical for optimal ensemble PRS performance and should be pursued whenever possible. We would also like to note that population mismatch is an issue not just for ensemble PRS training, but also for the general PRS training which has previously been discussed^23^.

### PUMAS-ensemble provides significant improvement over single PRS models

Lastly, we show that the performance improvement of our method is statistically meaningful. We compared the performance of PUMAS-ensemble with single PRS models across all analyzed traits and genetic ancestries (**Methods**). PUMAS-EN and PUMAS-SL demonstrated statistically significant improvement over the median PRS by 158.62% (p-value=5.7e-14; one-sided Wilcoxon signed-rank test) and 158.55% (p-value=5.7e-14), respectively (**Supplementary Table 16**). Moreover, we found that PUMAS-EN and PUMAS-SL also consistently outperformed the single best PRS in each analysis by an average of 11.47% (p-value=1.33e-06) and 11.27% (p-value=1.67e-06). We note that these gains were particularly striking in cross-ancestry applications (**Supplementary Table 16**). Overall, PUMAS-ensemble attains statistically significant improvement over fine-tuned single PRS method without accessing individual-level datasets.

## Discussion

Ensemble learning can effectively combine multiple PRS models and improve risk prediction accuracy, but it is a data-demanding task that is often impossible to implement in practice due to the lack of adequately large individual-level holdout datasets. In this study, we introduced two summary-statistics-based ensemble learning approaches based on elastic net and super learning under the PUMAS-ensemble framework. Our proposed approaches employ statistical regularization to allow adaptive integration of a large number of single PRS models that may be highly correlated. We demonstrate that PUMAS-ensemble PRS closely approximates the ensemble PRS trained based on individual-level holdout data and show its superior performance compared to single PRS models in both within-ancestry and cross-ancestry applications.

Our work brings several key advancements to the field. First, PUMAS-ensemble is the only method in the literature that performs PRS ensemble learning on GWAS summary statistics, bypassing the stringent data requirement in existing approaches and fully exploiting the widely available summary-level GWAS data resources without compromising predictive performance. Second, our approach employs sophisticated regularization, allowing researchers to combine possibly hundreds of PRS models without acquiring additional holdout samples. Importantly, this strategy can build ensemble scores for non-European ancestries by combining a variety of ancestry-specific PRS models. While cross-ancestry ensemble learning has proven effective in improving upon single PRS models across several recent studies, existing strategies^21-23,26,27^ rely on non-European data at the individual level which can be close to impossible to obtain in practice. Our approach removes this critical constraint in data requirement which is a major step towards reducing disparity in genomic medicine^53,55^. Third, PRS method development is a crowded research field. When a new PRS method is introduced, it is common to see incremental gains in predictive accuracy over existing approaches. Our method shows a consistent 10-30% *R*^2^ improvement in within-ancestry applications and an *R*^2^ gain of as high as 300% in cross-ancestry predictions. This is a substantial improvement. A school of thought against ensemble learning in the PRS research field is that although there is variation in the performance of different models, the best single PRS would perform similarly well with reduced computational burden compared to the ensemble score. We have demonstrated that our method achieves substantial and statistically significant gain over the fine-tuned single PRS. In addition, although it seems intuitive to simply pick the best-performing PRS for downstream application, even this (conventionally) requires a dataset with individual-level genotype and phenotype information for model evaluation. This is a key point we have emphasized throughout this paper––some basic tasks in PRS optimization can be data-demanding, which is why it is crucial to develop statistical methods that can achieve these tasks on summary statistics alone. Currently, in a highly realistic scenario where only one GWAS summary-level dataset is available, a summary-statistics-based cross-validation method (e.g., PUMAS^5,6^) will have to be employed for PRS model selection. Our study has introduced a better alternative to further improve the performance of the single best PRS model without introducing unrealistic data requirement or computational burden. Further, perhaps the most important feature of our approach is its ability to continuously evolve. We did not introduce just another PRS model. This is a highly adaptive framework that can always incorporate everyone’s favorite PRS models, including future models once they become available. This highlights summary statistics-based ensemble learning as a crucial direction for future PRS development, and is also why we believe we may have found the “one score to rule them all”. Every future PRS method should consider this strategy to combine the cumulative wisdom from many existing models with new methodological innovations. Summary-statistics-based ensemble learning is the core technique that makes this possible.

There are still some important future directions. In this study, we included several PRS methods for illustration. Additional models researchers can consider in their analysis include methods that introduce new statistical designs^30,56^, employ non-parametric modeling^57,58^, leverage multi-trait joint modeling^12-14,20,37,59^, or incorporate biological annotations^10,11,35^. It has been shown that ensemble learning of PRS trained on multiple traits can improve risk prediction performance^20^. Thus, multi-trait ensemble learning based on summary statistics would be a reasonable future extension of PUMAS-ensemble. However, this raises a more general question about how to properly select PRS input in PUMAS-ensemble. We highlight the flexibility of PRS input as a major feature of our method––PUMAS-ensemble can adaptively and efficiently integrate hundreds of scores and users can decide what PRS models they would like to include based on their goal and computational infrastructure. We recommend users to include PRS models generated from diverse statistical designs targeting distinct genetic architecture. Still, it remains an important future direction to develop a systematic strategy for input PRS selection. We also did not consider non-linear models for either PRS construction or ensemble learning^20,21,60^. This is related to the empirical observation that PUMAS-SL shows highly comparable performance to PUMAS-EN despite being conceptually more complex. Our results suggest that the added complexity of super learner might not yet yield substantial benefits under the current landscape of PRS methods where only linear models are combined, although PUMAS-SL may be worth a reassessment in the future if effective non-linear models are introduced in the field. Another limitation of this study is that we only benchmarked PRS performance using R^2^ and AUC. Recent studies have suggested other important measures of model performance which may be more relevant to specific PRS applications^61-63^. We recommend researchers to choose the most relevant metric based on their goals when applying PUMAS-ensemble. Due to the instability in risk estimates from different PRS methods, one specific recent PRS application is stable PRS over time for clinical use^62,63^. It has been shown that continually updating PRS using ensemble learning with individual-level data leads to significantly more stable PRS performance compared to choosing the singular model with the best predictive performance at the time^63^. A current limitation of these methods is that they require independent ensemble training dataset never used in any training exercise, including the new PRS being added, which may have new samples. PUMAS-ensemble can bring some flexibility to this limitation. If a new PRS model is published, one can add this model to PUMAS-ensemble using the same GWAS that other models were trained with. However, current implementations of these models ensemble weights derived from PGS catalog over time. PUMAS-ensemble will not work with black box PRS models where we do not know the input GWAS, such as models from PGS catalog. Robustness to sample overlap would be required to implement a similar strategy using summary statistics alone, and this represents an interesting extension of PUMAS-ensemble. Additionally, it would be meaningful work to extend all recent multi-ancestry PRS methods that use ensemble learning on holdout samples^21-23,26,27^ to the summary statistics-based version using PUMAS-ensemble. Finally, it remains an open question how population admixture^64^ and ancestry continuum^65^ should be modeled. In conclusion, our study presents a highly innovative and data-efficient statistical framework for PRS ensemble learning. We highlight its capability of combining, and thus surpassing, all existing (and future) PRS models. PUMAS-ensemble is a versatile tool that can bring immediate benefits to the many applications of PRS which will no doubt greatly facilitate future studies.

## Methods

### An overview of summary-statistics-based PRS ensemble learning

Conventionally, PRS ensemble learning requires a summary-level GWAS dataset for single PRS model training and an independent individual-level dataset for ensemble model training. If the ensemble model contains tuning parameters, such as regularization parameters in penalized regressions, the individual-level dataset needs to be first partitioned for model tuning and evaluation, and eventually combined again for fitting the best ensemble model (e.g., training-validation split or cross-validation). Such a procedure is straightforward when the required individual-level dataset exists. In practice, however, it is much more common that only GWAS summary statistics are available. Therefore, we extend a flexible summary-statistics-based cross-validation approach we previously introduced^32^ to train and evaluate elastic net and super learning ensemble PRS models using only GWAS summary statistics and LD reference data as inputs. We first provide an overview of the PUMAS-ensemble framework. Under an additive genetic model, the relationship between a trait *Y* and genotype ***X*** = (***X***_1_, … , ***X***_*p*_) can be quantified as:

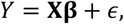

where *p* is the number of SNPs, ***β*** ∈ ℝ^*p*^ denote true SNP effects, and *ϵ* denotes distributed random error terms independent from ***X*** with zero mean and finite and positive variance *σ*^2^. We then define ensemble PRS as a linear combination of many PRS models:

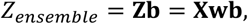

where ***b*** = [*b*_1_ *b*_2_ … *b*_*M*_]^*T*^ is the vector of ensemble weights for *M* PRS models, ***Z*** = [*Z*_1_ *Z*_2_ … *Z*_*M*_] = ***Xw*** is the PRS matrix of *M* models with corresponding SNP weights ***w*** = [***w***_1_ ***w***_2_ … ***w***_*M*_]. The key objective is to obtain the ensemble weights ***b*** for optimal PRS performance.

Assume that we have GWAS summary-level data of sample size *N* and external reference genotype data for LD estimation, in order to train and evaluate ensemble PRS models, we need to partition the full GWAS summary statistics into three subsets for PRS model training, ensemble learning, and benchmarking, respectively. We have shown in earlier work^31,32^ that given the observed summary-level data ***x***^*T*^***y*** from the full GWAS dataset, the conditional distribution of summary statistics for a subset of GWAS samples 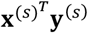 with sample size *N*^(*s*)^ < *N* is:

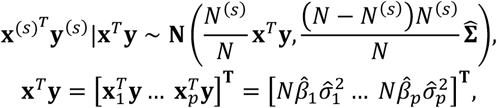

where 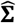 is the observed covariance matrix of ***x***^*T*^***y***, and 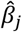 and 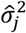 are the marginal effect size estimate and MAF-based variance estimator, respectively, for SNP *j, j* = 1,2, … , *p*. We have shown that 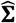 can be obtained based on GWAS summary statistics and LD reference data, and an iterative subsampling scheme can be used to partition the full summary statistics into three independent subsets of summary statistics^32^:

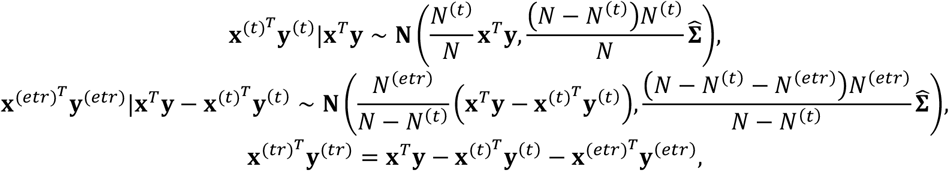

where 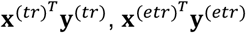, and 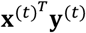 correspond to the summary statistics for PRS model training, ensemble learning, and benchmarking (testing), respectively. Details of PRS model fitting and training of ensemble models are described in later sections.

Once we obtain the ensemble weights ***b***, we can evaluate prediction accuracy of the ensemble model by calculating its predictive *R*^2^ on the testing summary-level data as^32^:

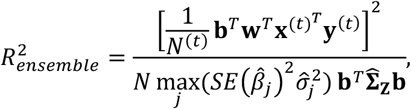

where 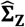 is the estimated covariance matrix of *M* PRS models, which can be approximated using the LD reference data. Finally, we repeat the above procedures *K* times (i.e., *K*-fold MCCV) to ensure robust performance of ensemble PRS. Note that if the goal is to evaluate ensemble PRS without accessing external, individual-level validation datasets, PUMAS-ensemble can report the average prediction accuracy of ensemble PRS as 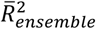across *K* folds. When the analytical aim is to produce ensemble PRS for maximal out-of-sample prediction accuracy, 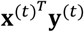 and 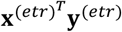 can be combined to calculate SNP weights for the ensemble model as 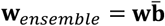, where 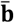 is the vector of average ensemble weights across *K* folds. In the following two sections, we will introduce the summary-statistics-based ensemble model fitting based on elastic net and super learning.

### PUMAS-EN: elastic net ensemble PRS based on summary statistics

Our earlier work has shown that combining multiple fine-tuned PRS models with linear regression can lead to better predictive performance than single PRS^32^. While classic linear regression serves as a proof-of-concept example for ensemble PRS, it is often of interest to aggregate as many PRS models as possible for achieving maximal gain in prediction accuracy. However, as demonstrated in our simulation study, summary-statistics-based linear regression becomes highly unstable and hinders performance of ensemble score when many highly correlated PRS models are included. To address multicollinearity and further improve ensemble score, we introduce PUMAS-EN which adaptively integrates a large number of PRS models via elastic net^66^ using GWAS summary statistics. Another advantage of this elastic net model is that it relieves PUMAS-ensemble from conducting fine-tuning for each PRS method prior to ensemble learning and can directly combine all PRS across various methods and tuning parameter settings.

PUMAS-EN has two tuning parameters, *λ* and *α*, where *λ* controls the overall shrinkage of each PRS’s coefficient in the ensemble model and *α* allocates the relative contribution of L1 and L2 penalty terms. To obtain elastic net coefficient estimates for ***b***, the ensemble weights for *M* single PRS models, PUMAS-EN minimizes the following objective function:

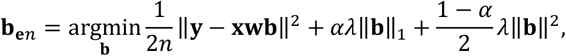

where *n* is sample size of elastic net training data, and ***xw*** are standardized PRS with mean of 0 and variance of 1. Following derivations from lassosum^4^ and recent PRS frameworks^12,22,23^ that utilized penalized regression, we show that ***b***_***e****n*_ can be estimated using only GWAS summary statistics and an external LD reference data. Since the objective function is not continuously convex, we use the coordinate descent^67^ algorithm to iteratively update ensemble weight for PRS model *m, m* = 1,2, … , *M*:

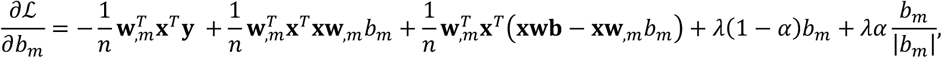

where ***w***_,*m*_ is the *m*-th column of ***w*** that represents SNP weights in the *m*-th PRS model. By setting 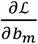 to zero, we obtain the formula for iteratively updating *b*_*m*_:

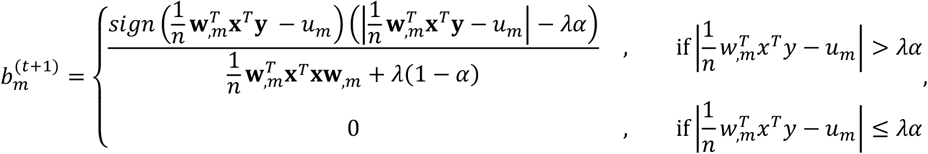

where 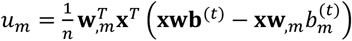P utilizes the PRS weights estimated from the *t*-thiteration 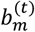.

It is clear that only the terms ***x***^*T*^***y*** and ***x***^*T*^***x*** are required for fitting elastic net ensemble model. To fine-tune hyperparameters *λ* and *α*, we further partition the ensemble training summary statistics 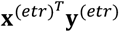 into two independent subsets. We fit multiple elastic net models with different combinations of *α* and *λ* on the first subset, select the “optimal” combination based on their performance on the second subset, and eventually train the fine-tuned elastic net model with the selected hyperparameter values on the entire ensemble training summary statistics 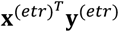. Throughout this study, we consider hyperparameter settings *α* = 0, 0.25, 0.5, 0.75, 1 and *λ* = 10ψ where ψ includes 51 numbers evenly spaced in [™5,0]. For model fitting, we initialize 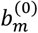 at zero and iteratively update 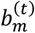 until the algorithm converges 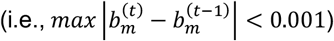 or when reaching the maximum number of iterations (default: 10^4^).

### PUMAS-SL: super learning PRS based on summary statistics

Super learning^24^ is essentially a two-level ensemble learning approach that trains an optimal weighted combination of multiple machine learning models such as elastic net^66^, ridge regression^68^, and LASSO^69^. It can be used to train an “ensemble of ensemble” model combining a large set of baseline PRS models for optimized polygenic risk prediction. All existing super learning strategies for PRS training require individual-level data as input. Here, we introduce PUMAS-SL, which trains a super learning model using only GWAS summary statistics.

First, we define super learning PRS as a linear combination of level-one ensemble PRS models:

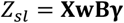

where ***B*** = [***b***_*en*_ ***b***_*ridge*_ ***b***_*LASS*0_] denotes the matrix of level-one ensemble weights, i.e., the weights of various PRS models in different ensemble models (elastic net, ridge regression, LASSO), and *γ* = [*γ*_*en*_ *γ*_*ridge*_ *γ*_*LASS*0_]^*T*^ denotes the level-two ensemble weights for the various ensemble models in the super learning PRS. Both ***B*** and *γ* are parameters of interests that need to be estimated. Using our subsampling scheme for summary statistics, we partition the ensemble training summary statistics 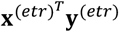 into two independent subsets denoted by 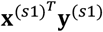 and 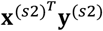to train ***B*** and *γ*, respectively. We fit the level-one ensemble models including ridge regression, LASSO, and elastic net on the first subset:

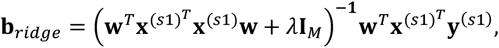

where ***b***_*LASS*0_ and ***b***_*en*_ can be estimated using the coordinate descent algorithm introduced earlier. To improve the stability of the super learning model, we further apply a *K*-fold summary-statistics-based MCCV on ^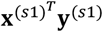^ and divide it into two independent subsets for training and fine-tuning level-one ensemble models, respectively. We then combine elastic net, ridge, and LASSO ensemble models through a level-two linear regression on ^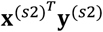^ from the second subset:

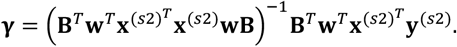

To ensure stable performance of PUMAS-SL, *γ* is estimated via non-negative least squares using coordinate descent algorithm. Taking ***B*** and *γ* together, we can obtain a super learning PRS model based only on GWAS summary statistics. For out-of-sample prediction, PUMAS-SL outputs SNP weights for super learning PRS model as 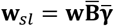, where 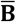 and 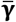 are estimated level-one and level-two ensemble weights averaged across *K*-fold MCCV.

### Details of PRS model training

We trained and combined PRS based on eight methods including lassosum^4^, LDpred2^6^, PRS-CS^5^, MegaPRS^8^, SBayesR/SBayesRC^7,35^, DBSLMM^9^, Vilma^33^, SBLUP^34^ throughout this study. Single PRS models were trained separately on each chromosome in parallel, except for the ones trained by MegaPRS and lassosum. We constructed LD reference data for PRS methods that do not provide such datasets throughout this study. For EUR (i.e., simulation, UKB, and AllofUs) and EAS PRS analyses (i.e., AllofUs), we used a UKB genotype dataset consisting of N=1000 randomly selected individuals of EUR ancestry and another UKB EAS genotype dataset (N=500) as the corresponding LD reference datasets, respectively. When training ancestry-specific PRS for blood lipid traits in UKB, we randomly picked 500 samples from the UKB testing dataset of matched ancestry to generate LD data for each of EUR, AFR, EAS, and SAS populations. We used HapMap 3 SNPs in all our analyses. For the rest of this section, we outline and briefly introduce each single PRS method considered in our study.

#### Lassosum

estimates LASSO coefficients by jointly modeling SNPs in LD using estimated marginal SNP effect sizes from GWAS summary statistics. Lassosum has two tuning parameters *s* and *λ*, where *s* regulates the sparsity of LD blocks and *λ* is the regularization parameter in LASSO that shrinks SNP effects towards zero. We trained lassosum models with *s* = 0.2, 0.5, 0.9 and *λ* = 0.005, 0.01 using the R package ‘lassosum’ (v0.4.5).

#### LDpred2

is a Bayesian PRS method that adaptively shrinks SNP effects while accounting for LD. LDpred2 employs two versions assuming two distinct prior distributions for SNP effects: LDpred2-inf, which is a tuning-free model based on the ‘infinitesimal model’, and LDpred2-grid, which assumes a spike-and-slab prior distribution with hyperparameters *p* representing the true proportion of causal variants and total heritability h^2^. In addition, LDpred2-auto is an empirical Bayes approach that avoids the need for hyperparameter tuning by estimating *p* and h^2^ along with other parameters during model fitting^6^. We included LDpred2-grid and LDpred2-auto models in ensemble PRS in all our analyses. We trained LDpred2 models using the R package ‘bigsnpr’ (v1.9.11) with *p* = 0.001, 0.01, 0.1 and low heritability 0.1 *⋅* h^2^, 0.3 *⋅* h^2^, where h^2^ is the heritability estimated by LD-score regression^70^, both sparse and non-sparse models for LDpred2-grid, and more stringent LD shrinkage (shrink_corr = 0.5) for LDpred2-auto. We adopted both lower heritability value and stronger LD shrinkage to improve LDpred2 model convergence following recent improvement made to LDpred2^71^.

#### PRS-CS

places a continuous shrinkage prior on SNP effect size distribution, as opposed to the spike-and-slab prior in LDpred2. It includes a global shrinkage factor *Φ* which uniformly shrinks SNP effects throughout the genome. Alternatively, *Φ* can be adaptively learned from the GWAS data by a fully Bayesian approach (PRS-CS-auto). We fitted PRS-CS models using 1000 Genomes Project EUR LD matrices in simulation and UKB LD matrices for real data analysis. All LD reference data were provided by the PRS-CS software (v1.0.0).

#### SBayesR

employs a mixture of point mass at zero and three normal distributions with different variance parameters as the prior distribution for SNP effects, representing SNP effect sizes of different magnitudes. SBayesR does not require hyperparameter tuning because all hyperparameter values are pre-specified. We fitted SBayesR models using the GCTB software (v2.04.3) and the sparse UKB LD matrices for HapMap 3 SNPs provided by GCTB. SBayesR was used in simulation study.

#### SBayesRC

extends SBayesR to incorporate functional annotation and improve LD modeling through a low-rank model. Importantly, SBayesRC addresses the model convergence issue that has been previously reported for SBayesR^35^. We trained the SBayesRC model using v0.2.6 of its software provided on GitHub, the provided annotation information from Baseline model 2.2, and generated custom LD reference data using provided instructions on GitHub for each analysis. We used SBayesRC instead of SBayesR in all real data analyses.

#### Vilma

is another recently developed Bayesian approach with a more flexible normal mixture prior than SBayesR. It can be applied to model summary statistics from multiple traits and different genetic ancestries. The number of component normal distributions in its mixture prior is the only tuning parameter in Vilma; like SBayesR, it recommends a default value (i.e., 81) for this parameter. We fitted Vilma models using its software provided on GitHub.

#### MegaPRS

is a flexible Bayesian framework that can employ multiple prior specifications such as LASSO, ridge, Bolt (i.e., a mixture of two Gaussian distributions), and BayesR (i.e., a mixture of three Gaussian distributions and a point mass). We fitted MegaPRS models using the LDAK software (v5.2) with the recommended BayesR prior specification which include 84 pairs of tuning parameters that determine the relative weights of component Gaussian distributions. LDAK-thin^72^ was used for per-predictor heritability estimation. To improve robustness of ensemble score, we only included MegaPRS models with no greater than 10 predictors that failed to converge.

#### DBSLMM

first conducts LD clumping and thresholding to partition SNPs into a large-effect group and a small-effect group. It then fits a linear mixed effects model (i.e., large fixed effects and small random effects) to obtain updated SNP weights using summary statistics while accounting for LD. DBSLMM is a computationally efficient approach that has one tuning parameter, the p-value threshold in LD clumping and thresholding. We fitted DBSLMM models using fine-tuned p-value cutoff determined by summary-statistics-based parameter tuning implemented in the DBSLMM software (v0.3).

#### SBLUP

bases its framework on a linear mixed model where SNP effects are assumed to be random and normally distributed, thus making it conceptually equivalently to LDpred-inf. SBLUP uses GWAS summary statistics as input and does not have hyperparameters. We trained SBLUP model using the GCTA software (v1.93.0).

### Simulation studies based on UKB genotype data

We conducted simulations using UKB genotype data imputed to the Haplotype Reference Consortium panel. We kept samples of European ancestry and removed genetic variants with MAF below 0.01, imputation *R*^2^ below 0.9, Hardy-Weinberg equilibrium test p-value below 1e-6, or missing genotype call rate greater than 2%. We further extracted variants in the HapMap 3 SNP list and the LD reference data for European ancestry from Phase 3 of the 1000 Genomes Project. 377,509 samples and 944,547 variants remained after quality control. Then, we randomly selected 100,000 samples to form the simulation dataset with their genotype and randomly selected 1,000 samples to form the LD reference dataset. To generate phenotypic values, we simulated SNP effect sizes from a spike-and-slab distribution, 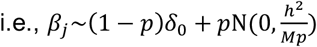, as assumed in LDpred2, where *p* is the proportion of causal variants, *δ*_O_ denotes a point mass at 0, h^2^ is the total heritability of the phenotype, and *M* is the total number of SNPs. We considered four simulation settings with distinct genetic architectures and heritability by setting *p* to 0.1% or 5% and h^2^ to 0.3 or 0.7. To simulate trait values, we randomly selected causal variants across the genome, computed the “true PRS” by aggregating SNP allele counts weighted by true effect sizes, and added gaussian noises scaled according to trait heritability. We fitted marginal linear regressions using PLINK^73^ to obtain GWAS summary statistics in each simulation setting.

We compared ensemble PRS constructed by PUMAS-ensemble and by 4-fold MCCV using individual-level data. To implement 4-fold MCCV, in each fold, we randomly selected 60% of the samples as the training dataset (N=60,000), 30% as the ensemble training dataset (N=30,000), and the remaining 10% as the testing dataset (N=10,000). We fitted GWAS and PRS models on the training data, calculated ensemble weights from elastic net and super learning on the ensemble training data, and finally, evaluated PRS model performance on the testing data. For elastic net, we divided ensemble training data into two halves and fitted elastic net models using the R package ‘glmnet’ (v4.1)^67^ with a prespecified set of tuning parameters on the first subset. Then we benchmarked the performance of each model on the second subset and re-trained the most predictive elastic net model on the entire ensemble training data. We used the R package ‘SuperLearner’ (v2.0)^24^ to train super learning PRS on the ensemble training dataset. As a comparison, we applied PUMAS-ensemble to implement 4-fold summary-statistics-based MCCV. We partitioned the full summary-level data into three independent sets of summary statistics for PRS training (N=60,000), ensemble learning (N=30,000), and PRS benchmarking (N=10,000), respectively. For PUMAS-EN, we trained fine-tuned ensemble model by dividing the ensemble training summary statistics into two halves and following the PUMAS-EN model fitting strategy we introduced earlier. For PUMAS-SL, we partitioned the ensemble training summary-level data to two subsets for training level-one (2/3; N=40,000) and level-two (1/3; N=20,000) ensemble weights, respectively. An additional 50%-50% training-testing split on the level-one ensemble training subset was applied to train fine-tuned LASSO, ridge, and elastic net regression models. To test if collinearity was driving the degradation of linear regression performance, we increased the number of PRS models by 50 and 100 correlated fake PRS weights for the simulation setting with h^2^ = 0.3 and *p* = 0.1%. These fake PRS weights were generated conditioning on a randomly selected true PRS model, assuming a bivariate normal distribution between the selected and fake PRS with a correlation of 0.5. Throughout simulation, we used European LD reference data from Phase 3 of the 1000 Genomes Project^74^ to subsample summary statistics and evaluate PRS performance. The holdout UKB LD genotype data (N=1,000) was used as the LD reference for PRS model training, except for PRS-CS^5^ and SBayesR^7^, where software-provided UKB LD matrices were used. Finally, for both approaches, we calculated and reported average *R*^2^ for each PRS model across 4 folds.

### Evaluating ensemble PRS in UKB

We compared the ensemble PRS constructed by PUMAS-ensemble with those constructed by training-testing split based on individual-level data for 16 quantitative phenotypes on UKB individuals of EUR descent. The list of UKB phenotypes and detailed sample size information are summarized in **Supplementary Table 3**. We reserved approximately 10% of the UKB samples with non-missing phenotypic values (N=38,521) as the holdout set for individual-level ensemble PRS training and PRS benchmarking. We then obtain GWAS summary statistics for each trait by performing linear regression analysis adjusting for sex, age polynomials to the power of two, interactions between sex and age polynomials, and top 20 genetic principal components via Hail (v0.2.57)^75^ on the remaining samples. A smaller subset of the holdout data (N=1,000) was used as LD reference for PRS model training. Prior to evaluating each PRS model, we regressed out covariate effects from phenotypes in the holdout dataset. Predictive *R*^2^ was reported for each PRS model.

To fit ensemble PRS based on individual-level data, we randomly partitioned the holdout dataset into two subsets for ensemble model training (3/4 of samples) and PRS benchmarking (1/4 of samples), respectively. We trained single PRS methods based on full GWAS summary statistics, fitted elastic net and super learning PRS on the ensemble training subset, and calculated predictive *R*^2^ for all PRS on the benchmarking subset. We trained fine-tuned elastic net model, super learning model, and PRSmix ^37,63^ on the ensemble training subset following the same protocols described in our simulation study but without cross-validation. We trained PRSmix based on default parameters (power threshold = 0.95; p-value threshold = 0.05). For comparison, we used PUMAS-ensemble to build ensemble PRS models by 4-fold MCCV. Within each fold of MCCV, we partitioned full GWAS summary statistics into training (70%) and ensemble training (30%) summary statistics. Model fitting for PUMAS-EN and PUMAS-SL follows the exact procedures described in PUMAS-ensemble simulation. Then, we computed and evaluated PUMAS-ensemble PRS on the benchmarking subset. The same subset of UKB holdout data (N=1,000) was used for subsampling summary statistics and ensemble model training for PUMAS-ensemble. Finally, we sought to compare PUMAS-ensemble with conventional ensemble learning under a common scenario where the individual-level data for ensemble model training is limited. To mimic this real-world setting, we randomly selected 500 samples from the ensemble model training subset to train elastic net and super learning ensemble PRS models and assessed their performance on the same PRS benchmarking subset. We also compared PUMAS-ensemble with PRS-PCA^38^ based on the same set of input PRS models. PRS-PCA was obtained as the first principal component of the PRS matrix for the testing data.

To benchmark the computational cost and scalability of the ensemble step of our approach, we performed simulation using the height GWAS data. Based on a PRS weights matrix with approximately 1.2 million SNPs and 51 PRS models, we gradually increased the number of PRS models to 100, 200, and 500 by simulating fake PRS weights. These fake PRS weights were generated conditioning on a randomly selected true PRS model, assuming a bivariate normal distribution between the selected and fake PRS with a correlation of 0.5. Runtime (based on 4 CPU threads) and memory usage was reported.

### Extracting phenotypes in AllofUs

We evaluated PRS performance for ten phenotypes in AllofUs: standing height, body mass index, diastolic blood pressure, systolic blood pressure, bipolar disorder, schizophrenia, major depressive disorder, type 2 diabetes, coronary artery disease, and breast cancer. Measures for height, body weight, diastolic and systolic blood pressure were obtained from the Physical Measurements in AllofUs using “measurement_concept_id” values 3036277, 3025315, 3012888, and 3004249. We required standing height and weight measurements to have units of centimeters and kilograms, respectively. body mass index was calculated as weight (kg) divided by height squared (m^2^). Following the AllofUs blood pressure protocol, only participants with three repeated blood pressure measurements were retained; we excluded the first measurement and averaged the second and third to derive diastolic and systolic blood pressure. For standing height, body mass index, and blood pressure, observations lying outside ±1.5 times the interquartile range were treated as outliers and removed. Disease phenotypes were defined from electronic health record data using “condition_concept_id” values (**Supplementary Table 7**). For cases, we used the earliest recorded condition date to compute age at diagnosis, while for controls, age was computed using the date of electronic health record consent (observation_concept_id = 1586099) for downstream regression analyses. Finally, we used genetic ancestry inferred from genetic principal components and sample relatedness provided in AllofUs to identify unrelated individuals within each ancestral group. We restricted the sample to individuals with biological sex of male or female in the analyses.

### External validation of PUMAS-ensemble PRS in AllofUs EUR samples

We compared PUMAS-EN and PUMAS-SL with single PRS models on independent samples of European descent in AllofUs^47^. We trained ensemble PRS models using publicly available GWAS summary statistics for standing height, body mass index, and six disease outcomes. Information about these GWAS datasets is summarized in **Supplementary Table 6**. For binary traits, we transformed logistic regression association statistics to linear scale before subsampling summary statistics^31^. For each trait, we partitioned the full summary statistics into two subsets for PRS training (70%) and ensemble model fitting (30%), respectively. SNP weights for elastic net and super learning PRS were obtained following the same analytical procedures used in our simulation study. We computed PRS for AllofUs samples using genotype data for HapMap 3 SNPs extracted from WGS data via PLINK1.9^73^. To evaluate PRS, we computed *R*^2^ for height and body mass index in AllofUs. Covariates including biological sex, age polynomials to the power of two, interactions between sex and age polynomials, and top 16 genetic principal components. We reported AUC as the performance metric for 6 disease endpoints. AUC values were computed from logistic regression between disease status and PRS with and without considering covariates. F-test based on these two models was performed to determine if ensemble PRS significantly improved model fit on top of using only covariates for disease prediction. To compute baseline-adjusted AUC improvement, we divided the AUC difference between PUMAS and the median single-PRS method by the median method’s AUC minus 0.5, to account for the percentage improvement, normalizing improvements relative to the signal above random guessing. We also trained and benchmarked PRSmix^37,63^ based on default parameters and input PRS models used to train PUMAS-ensemble. We divided the holdout dataset for each ancestry in two halves for training and benchmarking PRSmix against PUMAS-ensemble, respectively. Analysis of breast cancer PRS was restricted to females only. All PRS calculation and evaluation were conducted in the AllofUs cloud analysis environment using the v8 data release.

### Cross-ancestry ensemble PRS for EAS samples in AllofUs

We benchmarked PUMAS-ensemble against single PRS models based on GWAS summary statistics of European and East Asian ancestral populations, respectively, on East Asian samples in AllofUs^47^. We considered four continuous traits including height, body mass index, diastolic blood pressure, and systolic blood pressure and used independent EAS samples for PRS evaluation. Detailed data information is summarized in **Supplementary Table 10**.

We trained PUMAS-EN and PUMAS-SL using published EAS GWAS from BBJ^48-50^ and in-house EUR GWAS from UKB^32^. Sample size for each BBJ and UKB GWAS summary-level dataset is summarized in **Supplementary Table 10**. We fitted EUR PRS models using various PRS methods based on the full UKB GWAS summary statistics. Then, we applied PUMAS-ensemble to partition full BBJ summary statistics into two subsets for EAS PRS training (70%) and ensemble model fitting (30%), respectively. We used the same UKB EUR LD reference data (N=1,000) in UKB data analysis for EUR PRS training and a random subset of UKB EAS samples (N=500) for EAS PRS model fitting. The assignment of EAS ancestry for non-European UKB participants was described in our earlier work^26^. SNP weights for PUMAS-EN and PUMAS-SL were obtained based on the same procedures outlined in our simulation study. PRSmix was trained and benchmarked as described in the previous section.

PRS-CSx was trained using the default hyperparameters (a=1, b=0.5) and the auto flag which learns the global shrinkage parameter automatically from the data using a full Bayesian approach^27^. Since PUMAS-ensemble only uses summary statistics for prediction, we used ancestry-specific PRS-CSx auto results without validation data for a fair comparison. PRS-CSx results were not included in the ensemble learning step for PUMAS-EN or PUMAS-SL due to additional computational burden this would have required, however, doing so likely would have improved PUMAS-ensemble prediction.

We computed and evaluated EAS PRS, EUR PRS, PRS-CSx, PUMAS-EN PRS, and PUMAS-SL PRS models on the testing dataset. The same set of covariates considered in the AllofUs data analysis in the previous section have also been adjusted for prior to computing PRS *R*^2^.

### Cross-ancestry ensemble PRS of blood lipid traits for four ancestries in UKB

We compared PUMAS-ensemble against single PRS models based on GWAS summary statistics from GLGC (UKB samples excluded) of AFR, EAS, EUR, and SAS ancestral populations^51^ on independent samples in UKB. We considered four blood lipid traits HDL cholesterol, LDL cholesterol, logTG, and TC and used independent UKB samples with genetically predicted AFR, EAS, EUR, and SAS ancestries^22^. GLGC additionally provides GWAS summary statistics of AMR ancestry, which we excluded due to our standard QC pipeline flagging sample size issues on chromosomes 13 through 22 for both LDL cholesterol and logTG traits. An additional reason for excluding AMR samples was their limited samples in the independent UKB data.

For each ancestry and trait pair, we trained PUMAS-EN and PUMAS-SL using ancestry-specific GLGC summary statistics. Sample size for the GLGC summary statistics for each ancestry and trait pair is available in **Supplementary Table 12**. For each ancestry and trait, we applied PUMAS-ensemble to partition full summary statistics into two subsets for PRS training (70%) and ensemble model fitting (30%). We fitted single PRS models for the given trait for all ancestries using the same PRS methods described in previous sections which we later include in the ensemble learning. We trained PRS-CSx for each trait and ancestry as was described in the EAS cross-ancestry section. We used UKB LD reference data from a matching ancestry for each ancestry PRS training and model fitting. PUMAS-EN and PUMAS-SL SNP weights were obtained with similar procedures as the simulation study. Covariates including biological sex, age polynomials to the power of two, interactions between sex and age polynomials, and the top 10 genetic principal components were regressed out from both the phenotype and PRS. PRSmix was trained and benchmarked as described in the previous sections.

We further investigate the effect of LD misspecifications on PUMAS-ensemble in cross-ancestry genetic prediction. During the ensemble training step of PUMAS-ensemble, we re-trained PUMAS-ensemble models using 1000 Genomes Project LD data instead of the UKB LD reference while keeping everything else unchanged. We explored two scenarios of LD mismatch. First, we compared external (1KG) and target-sample (UKB) that are from the same ancestral background as the testing dataset. Second, for external LD panels (1KG), we evaluated the loss in prediction accuracy when moving from ancestry-matched LD data to mis-specified LD (e.g., using 1KG EAS LD instead of 1KG EUR LD for prediction in UKB EUR samples).

### Significance test for the comparison between PUMAS-ensemble and single PRS models

We conducted statistical tests to quantify whether improvement from PUMAS-ensemble comparing to median-/best-performing PRS is indeed statistically significant. For each real data analysis, we divided the individual-level holdout dataset into two halves, selected the median-/best-performing PRS on the first half, and benchmarked its performance with PUMAS-EN/PUMAS-SL on the second half. Taken all real data analyses together, we performed a one-sided Wilcoxon signed-rank test with the null hypothesis that percentage improvement from the selected PRS to PUMAS-ensemble is zero. Statistical tests were conducted separately for PUMAS-EN and PUMAS-SL, and for median-performing and best-performing PRS models.

## Code availability

PUMAS-ensemble software is freely available at https://github.com/qlu-lab/PUMAS.

## Author Contributions

Z.Z. and Q.L. conceived and designed the study.

Z.Z. developed the statistical framework.

Z.Z. and S.D. performed the statistical analysis. Y.W., S.D., and A.B. performed AllofUs data analysis.

X.Y. assisted in UKB simulation analysis.

Z.Z. and S.D. implemented the software.

Q.L. and J.J. advised on statistical and genetic issues. Z.Z., S.D., and Q.L. wrote the manuscript.

All authors contributed to manuscript editing and approved the manuscript.

## Acknowledgment

The authors gratefully acknowledge research support from National Institutes of Health (NIH) grant R21 AG085162, and support from the University of Wisconsin-Madison Office of the Chancellor and the Vice Chancellor for Research and Graduate Education with funding from the Wisconsin Alumni Research Foundation (WARF). This research has been conducted using the UK Biobank Resource under Applications 42148 and 17731. This study makes use of summary statistics from GWAS consortia. We thank GWAS investigators for providing publicly accessible GWAS summary statistics. This research uses data from AllofUs. The All of Us Research Program is supported by the National Institutes of Health, Office of the Director: Regional Medical Centers: 1 OT2 OD026549; 1 OT2 OD026554; 1 OT2 OD026557; 1 OT2 OD026556; 1 OT2 OD026550; 1 OT2 OD 026552; 1 OT2 OD026553; 1 OT2 OD026548; 1 OT2 OD026551; 1 OT2 OD026555; IAA #: AOD 16037; Federally Qualified Health Centers: HHSN 263201600085U; Data and Research Center: 5 U2C OD023196; Biobank: 1 U24 OD023121; The Participant Center: U24 OD023176; Participant Technology Systems Center: 1 U24 OD023163; Communications and Engagement: 3 OT2 OD023205; 3 OT2 OD023206; and Community Partners: 1 OT2 OD025277; 3 OT2 OD025315; 1 OT2 OD025337; 1 OT2 OD025276. In addition, the All of Us Research Program would not be possible without the partnership of its participants.

## References

1. Yang, J., et al. Common SNPs explain a large proportion of the heritability for human height. Nat Genet 42, 565–569 (2010).

2. International Schizophrenia, C., et al. Common polygenic variation contributes to risk of schizophrenia and bipolar disorder. Nature 460, 748–752 (2009).

3. Vilhjálmsson, B.J., et al. Modeling Linkage Disequilibrium Increases Accuracy of Polygenic Risk Scores. Am J Hum Genet 97, 576–592 (2015).

4. Mak, T.S.H., Porsch, R.M., Choi, S.W., Zhou, X. & Sham, P.C. Polygenic scores via penalized regression on summary statistics. Genet Epidemiol 41, 469–480 (2017).

5. Ge, T., Chen, C.Y., Ni, Y., Feng, Y.A. & Smoller, J.W. Polygenic prediction via Bayesian regression and continuous shrinkage priors. Nat Commun 10, 1776 (2019).

6. Privé, F., Arbel, J. & Vilhjálmsson, B.J. LDpred2: better, faster, stronger. Bioinformatics 36, 5424–5431 (2020).

7. Lloyd-Jones, L.R., et al. Improved polygenic prediction by Bayesian multiple regression on summary statistics. Nat Commun 10, 5086 (2019).

8. Zhang, Q., Privé, F., Vilhjálmsson, B. & Speed, D. Improved genetic prediction of complex traits from individual-level data or summary statistics. Nature Communications 12, 4192 (2021).

9. Yang, S. & Zhou, X. Accurate and Scalable Construction of Polygenic Scores in Large Biobank Data Sets. Am J Hum Genet 106, 679–693 (2020).

10. Hu, Y., et al. Leveraging functional annotations in genetic risk prediction for human complex diseases. PLoS Comput Biol 13, e1005589 (2017).

11. Márquez-Luna, C., et al. Incorporating functional priors improves polygenic prediction accuracy in UK Biobank and 23andMe data sets. Nature Communications 12, 6052 (2021).

12. Chen, T.-H., Chatterjee, N., Landi, M.T. & Shi, J. A penalized regression framework for building polygenic risk models based on summary statistics from genome-wide association studies and incorporating external information. null, 1–19 (2020).

13. Hu, Y., et al. Joint modeling of genetically correlated diseases and functional annotations increases accuracy of polygenic risk prediction. PLoS Genet 13, e1006836 (2017).

14. Maier, R.M., et al. Improving genetic prediction by leveraging genetic correlations among human diseases and traits. Nat Commun 9, 989 (2018).

15. Monti, R., et al. Evaluation of polygenic scoring methods in five biobanks shows larger variation between biobanks than methods and finds benefits of ensemble learning. Am J Hum Genet 111, 1431–1447 (2024).

16. Wang, Y., Tsuo, K., Kanai, M., Neale, B.M. & Martin, A.R. Challenges and Opportunities for Developing More Generalizable Polygenic Risk Scores. Annu Rev Biomed Data Sci 5, 293–320 (2022).

17. Ni, G., et al. A Comparison of Ten Polygenic Score Methods for Psychiatric Disorders Applied Across Multiple Cohorts. Biol Psychiatry 90, 611–620 (2021).

18. Pain, O., et al. Evaluation of polygenic prediction methodology within a reference-standardized framework. PLOS Genetics 17, e1009021 (2021).

19. Yang, S. & Zhou, X. PGS-server: accuracy, robustness and transferability of polygenic score methods for biobank scale studies. Brief Bioinform 23(2022).

20. Albiñana, C., et al. Multi-PGS enhances polygenic prediction by combining 937 polygenic scores. Nature Communications 14, 4702 (2023).

21. Zhang, H., et al. A new method for multiancestry polygenic prediction improves performance across diverse populations. Nature Genetics 55, 1757–1768 (2023).

22. Zhang, J., et al. An ensemble penalized regression method for multi-ancestry polygenic risk prediction. Nature Communications 15, 3238 (2024).

23. Jin, J., et al. MUSSEL: Enhanced Bayesian polygenic risk prediction leveraging information across multiple ancestry groups. Cell Genomics 4(2024).

24. van der Laan, M.J., Polley, E.C. & Hubbard, A.E. Super learner. Stat Appl Genet Mol Biol 6, Article25 (2007).

25. Naimi, A.I. & Balzer, L.B. Stacked generalization: an introduction to super learning. Eur J Epidemiol 33, 459–464 (2018).

26. Miao, J., et al. Quantifying portable genetic effects and improving cross-ancestry genetic prediction with GWAS summary statistics. Nat Commun 14, 832 (2023).

27. Ruan, Y., et al. Improving polygenic prediction in ancestrally diverse populations. Nature Genetics 54, 573–580 (2022).

28. Patel, A.P., et al. A multi-ancestry polygenic risk score improves risk prediction for coronary artery disease. Nat Med 29, 1793–1803 (2023).

29. Willer, C.J., Li, Y. & Abecasis, G.R. METAL: fast and efficient meta-analysis of genomewide association scans. Bioinformatics 26, 2190–2191 (2010).

30. Chen, T., Zhang, H., Mazumder, R. & Lin, X. Fast and scalable ensemble learning method for versatile polygenic risk prediction. Proceedings of the National Academy of Sciences 121, e2403210121 (2024).

31. Zhao, Z., et al. PUMAS: fine-tuning polygenic risk scores with GWAS summary statistics. Genome Biol 22, 257 (2021).

32. Zhao, Z., et al. Optimizing and benchmarking polygenic risk scores with GWAS summary statistics. Genome Biology 25, 260 (2024).

33. Spence, J.P., Sinnott-Armstrong, N., Assimes, T.L. & Pritchard, J.K. A flexible modeling and inference framework for estimating variant effect sizes from GWAS summary statistics. bioRxiv, 2022.2004.2018.488696 (2022).

34. Robinson, M.R., et al. Genetic evidence of assortative mating in humans. Nature Human Behaviour 1, 0016 (2017).

35. Zheng, Z., et al. Leveraging functional genomic annotations and genome coverage to improve polygenic prediction of complex traits within and between ancestries. Nature Genetics 56, 767–777 (2024).

36. Bycroft, C., et al. The UK Biobank resource with deep phenotyping and genomic data. Nature 562, 203–209 (2018).

37. Truong, B., et al. Integrative polygenic risk score improves the prediction accuracy of complex traits and diseases. Cell Genomics 4(2024).

38. Coombes, B.J., Ploner, A., Bergen, S.E. & Biernacka, J.M. A principal component approach to improve association testing with polygenic risk scores. Genet Epidemiol 44, 676–686 (2020).

39. Zhang, H., et al. Genome-wide association study identifies 32 novel breast cancer susceptibility loci from overall and subtype-specific analyses. Nature genetics 52, 572–581 (2020).

40. Yengo, L., et al. A saturated map of common genetic variants associated with human height. Nature (2022).

41. Yengo, L., et al. Meta-analysis of genome-wide association studies for height and body mass index in approximately 700000 individuals of European ancestry. Hum Mol Genet 27, 3641–3649 (2018).

42. Trubetskoy, V., et al. Mapping genomic loci implicates genes and synaptic biology in schizophrenia. Nature 604, 502–508 (2022).

43. O’Connell, K.S., et al. Genomics yields biological and phenotypic insights into bipolar disorder. Nature 639, 968–975 (2025).

44. Adams, M.J., et al. Trans-ancestry genome-wide study of depression identifies 697 associations implicating cell types and pharmacotherapies. Cell 188, 640–652.e649 (2025).

45. Suzuki, K., et al. Genetic drivers of heterogeneity in type 2 diabetes pathophysiology. Nature 627, 347–357 (2024).

46. Aragam, K.G., et al. Discovery and systematic characterization of risk variants and genes for coronary artery disease in over a million participants. Nature Genetics 54, 1803–1815 (2022).

47. Bick, A.G., et al. Genomic data in the All of Us Research Program. Nature 627, 340–346 (2024).

48. Kanai, M., et al. Genetic analysis of quantitative traits in the Japanese population links cell types to complex human diseases. Nature Genetics 50, 390–400 (2018).

49. Akiyama, M., et al. Genome-wide association study identifies 112 new loci for body mass index in the Japanese population. Nature Genetics 49, 1458–1467 (2017).

50. Akiyama, M., et al. Characterizing rare and low-frequency height-associated variants in the Japanese population. Nature Communications 10, 4393 (2019).

51. Graham, S.E., et al. The power of genetic diversity in genome-wide association studies of lipids. Nature 600, 675–679 (2021).

52. Kachuri, L., et al. Principles and methods for transferring polygenic risk scores across global populations. Nat Rev Genet 25, 8–25 (2024).

53. Martin, A.R., et al. Clinical use of current polygenic risk scores may exacerbate health disparities. Nature Genetics 51, 584–591 (2019).

54. Hoggart, C.J., et al. BridgePRS leverages shared genetic effects across ancestries to increase polygenic risk score portability. Nature Genetics 56, 180–186 (2024).

55. Martin, A.R., et al. Human Demographic History Impacts Genetic Risk Prediction across Diverse Populations. Am J Hum Genet 100, 635–649 (2017).

56. Dun, Y., Chatterjee, N., Jin, J. & Nishimura, A. A Robust Bayesian Method for Building Polygenic Risk Scores using Projected Summary Statistics and Bridge Prior. arXiv preprint arXiv:2401.15014 (2024).

57. Zhou, G. & Zhao, H. A fast and robust Bayesian nonparametric method for prediction of complex traits using summary statistics. PLoS Genet 17, e1009697 (2021).

58. Chun, S., et al. Non-parametric Polygenic Risk Prediction via Partitioned GWAS Summary Statistics. Am J Hum Genet 107, 46–59 (2020).

59. Xu, C., Ganesh, S.K. & Zhou, X. mtPGS: Leverage multiple correlated traits for accurate polygenic score construction. The American Journal of Human Genetics 110, 1673–1689 (2023).

60. Elgart, M., et al. Non-linear machine learning models incorporating SNPs and PRS improve polygenic prediction in diverse human populations. Communications Biology 5, 856 (2022).

61. Wand, H., et al. Improving reporting standards for polygenic scores in risk prediction studies. Nature 591, 211–219 (2021).

62. Abramowitz, S.A., et al. Evaluating Performance and Agreement of Coronary Heart Disease Polygenic Risk Scores. Jama 333, 60–70 (2025).

63. Misra, A., et al. Instability of high polygenic risk classification and mitigation by integrative scoring. Nature Communications 16, 1584 (2025).

64. Sun, Q., et al. Improving polygenic risk prediction in admixed populations by explicitly modeling ancestral-differential effects via GAUDI. Nature communications 15, 1016 (2024).

65. Ruan, Y., et al. Leveraging genetic ancestry continuum information to interpolate PRS for admixed populations. medRxiv, 2024.2011. 2009.24316996 (2024).

66. Zou, H. & Hastie, T. Regularization and Variable Selection via the Elastic Net. Journal of the Royal Statistical Society. Series B (Statistical Methodology) 67, 301–320 (2005).

67. Friedman, J.H., Hastie, T. & Tibshirani, R. Regularization Paths for Generalized Linear Models via Coordinate Descent. Journal of Statistical Software 33, 1 – 22 (2010).

68. Hoerl, A.E. & Kennard, R.W. Ridge Regression: Biased Estimation for Nonorthogonal Problems. Technometrics 12, 55–67 (1970).

69. Tibshirani, R. Regression Shrinkage and Selection via the Lasso. Journal of the Royal Statistical Society. Series B (Methodological) 58, 267–288 (1996).

70. Bulik-Sullivan, B.K., et al. LD Score regression distinguishes confounding from polygenicity in genome-wide association studies. Nat Genet 47, 291–295 (2015).

71. Privé, F., Arbel, J., Aschard, H. & Vilhjálmsson, B.J. Identifying and correcting for misspecifications in GWAS summary statistics and polygenic scores. Human Genetics and Genomics Advances 3, 100136 (2022).

72. Speed, D., Hemani, G., Johnson, M.R. & Balding, D.J. Improved heritability estimation from genome-wide SNPs. American journal of human genetics 91, 1011–1021 (2012).

73. Purcell, S., et al. PLINK: a tool set for whole-genome association and population-based linkage analyses. American journal of human genetics 81, 559–575 (2007).

74. McVean, G.A., et al. An integrated map of genetic variation from 1,092 human genomes. Nature 491, 56–65 (2012).

75. Poterba, T., et al. The Scalable Variant Call Representation: Enabling Genetic Analysis Beyond One Million Genomes. bioRxiv, 2024.2001.2009.574205 (2024).

