## Supplementary Figures for "One score to rule them all: regularized ensemble polygenic risk prediction with GWAS summary statistics"

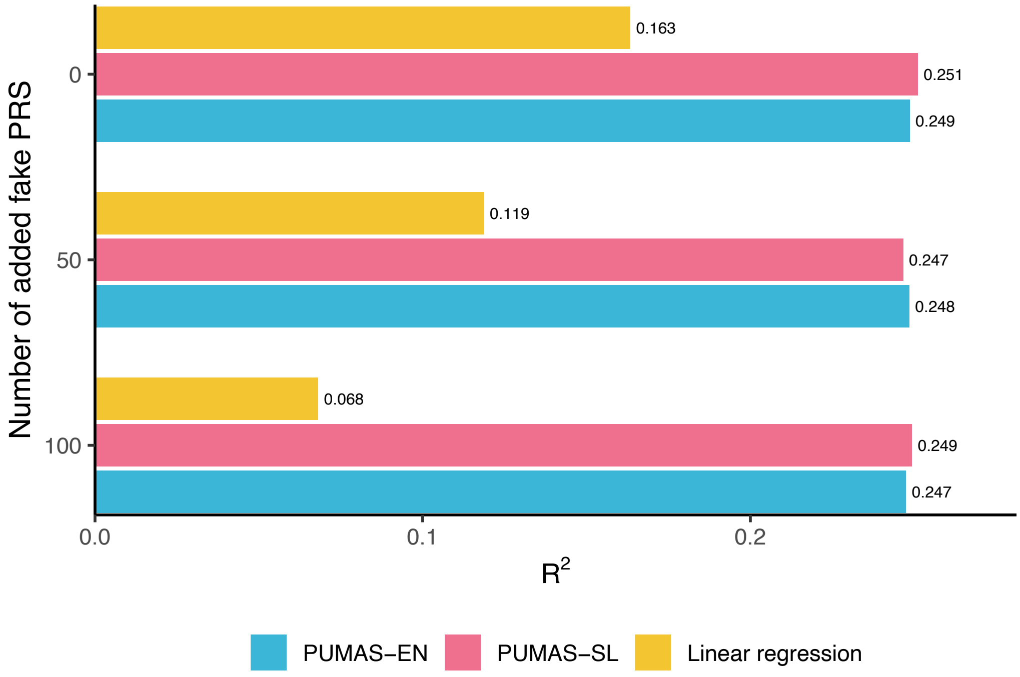


**Fig S1**. **PUMAS-ensemble performance with additional correlated PRS models.** We added 50 and 100 "fake" PRS models comprised of random noise and evaluate performance of ensemble models in the simulation setting with $h^2$ = 0.3 and $p$ = 0.1%.

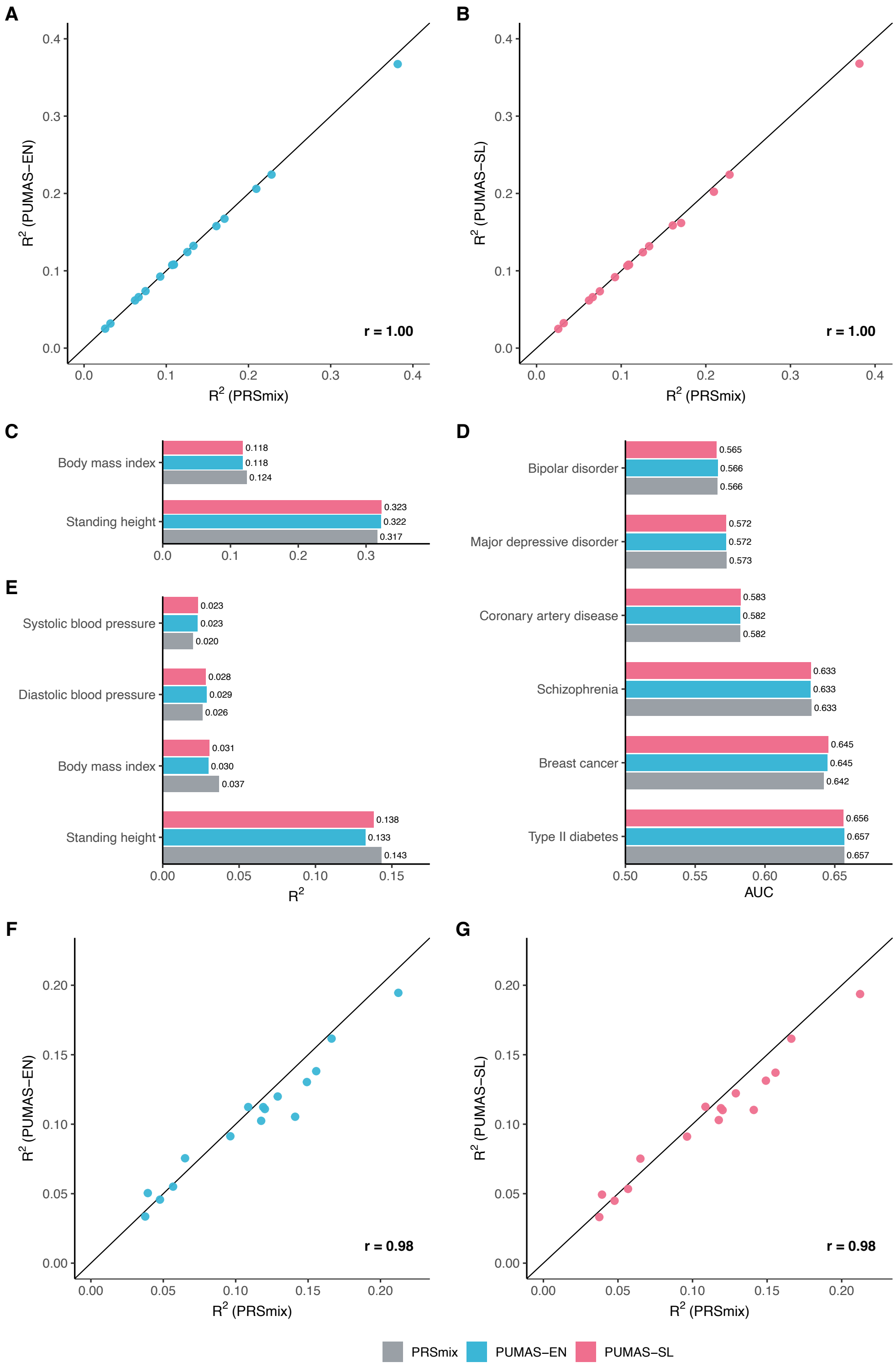


**Fig S2**. **Comparison between PRSmix and PUMAS-ensemble**. Both **(A)** PUMAS-EN and **(B)** PUMAS-SL were benchmarked with PRSmix trained from the same input PRS models for 16 traits in the UKB EUR dataset. Comparison between PRSmix and PUMAS-ensemble for (**C**) 2 continuous and (**D**) 6 binary traits trained using publicly available European GWAS, $R^2$ and AUC values from validation in AllofUs European cohort. Comparison between PRSmix and PUMAS-ensemble trained using publicly available East Asian and European GWAS evaluated in (**E**) East Asian participants in AllofUs. **(F)** PUMAS-EN and **(G)** PUMAS-SL were benchmarked with PRSmix fitted from the same input PRS models trained using GLGC GWAS sumstats for 4 lipid traits across 4 ancestries (AFR, EUR, EAS, SAS).

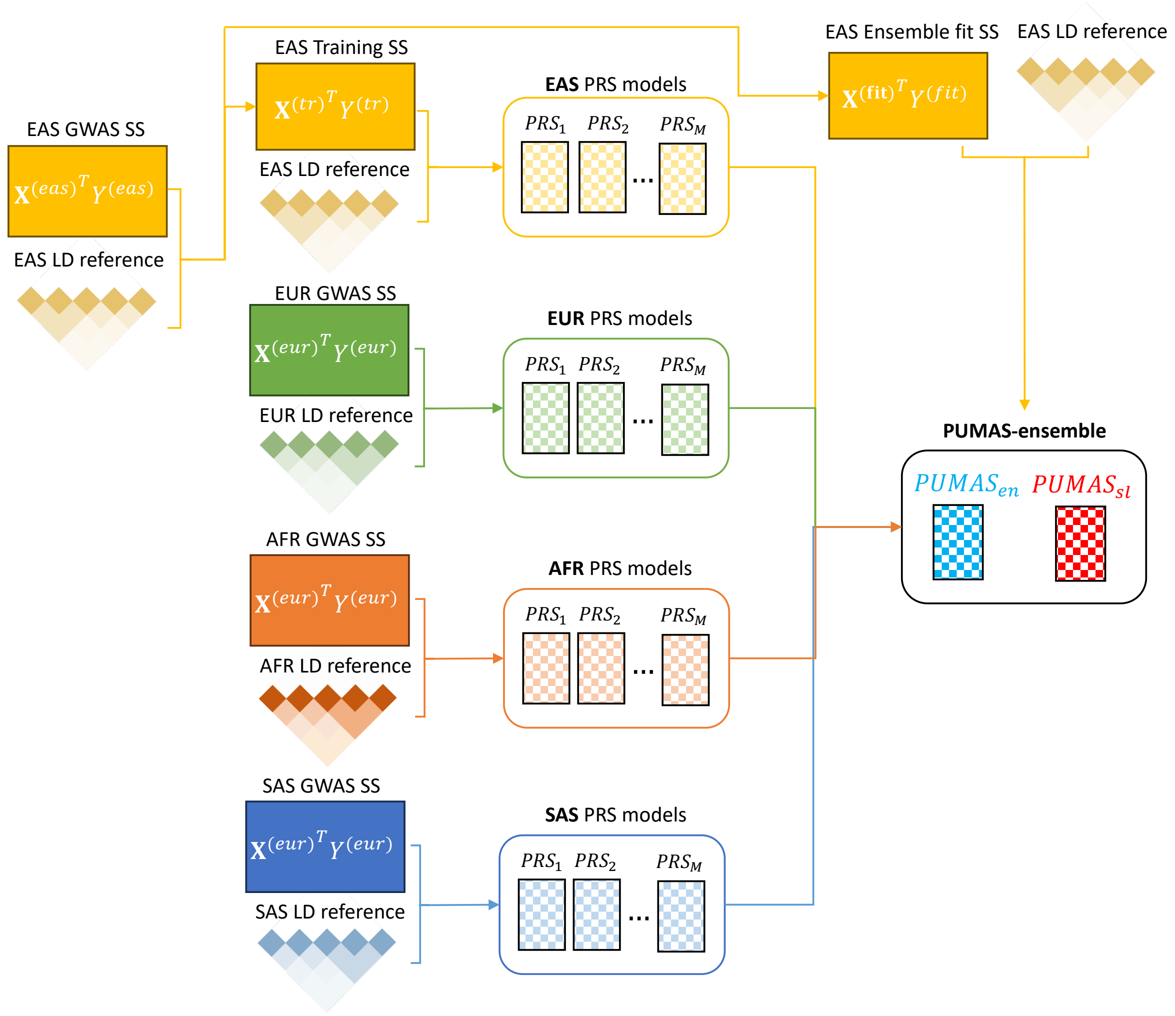


**Fig S3. Workflow for cross-ancestry PUMAS-ensemble application.** For illustration, we use EAS as the example for target ancestry.

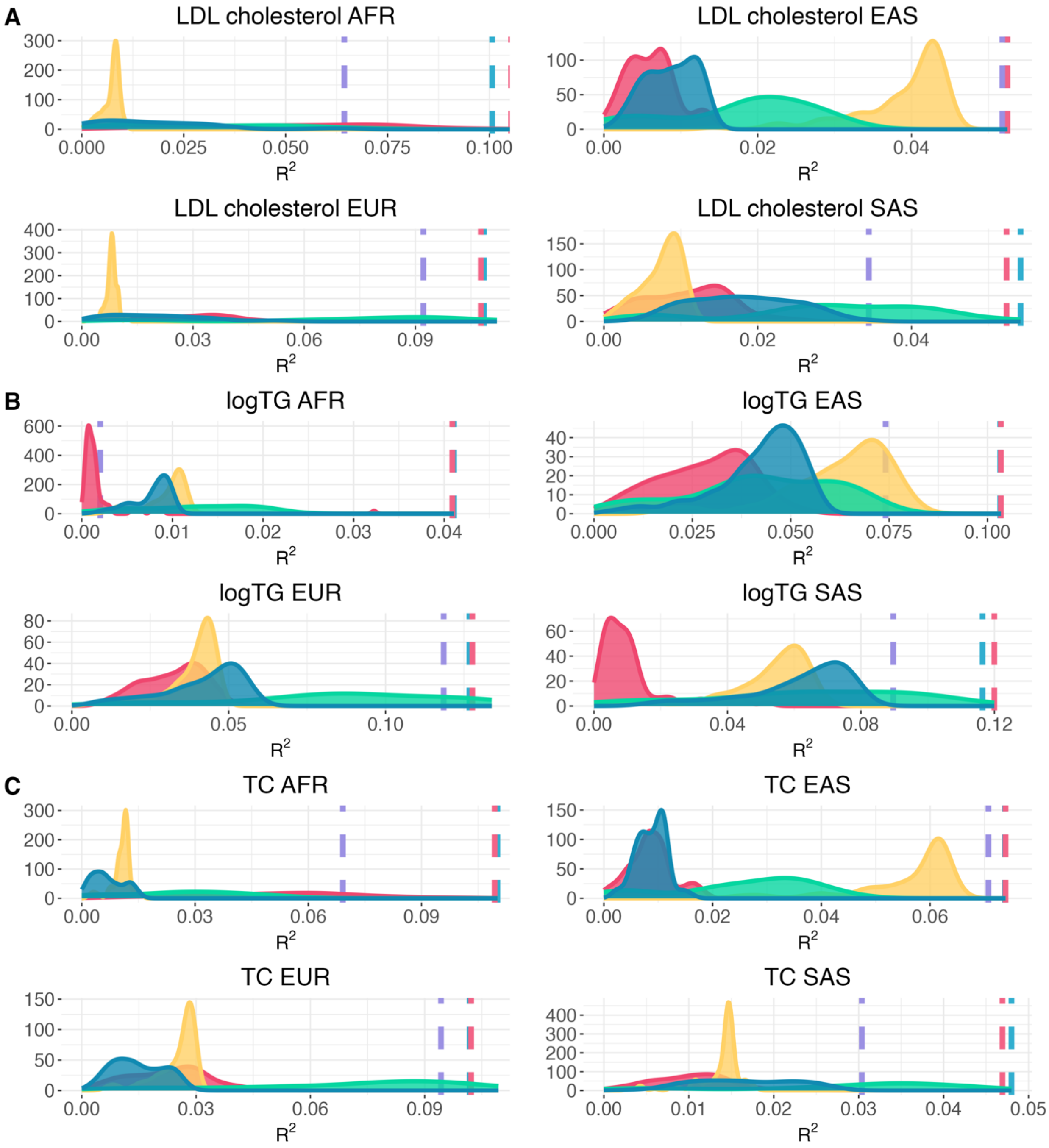


**Fig S4. Cross-ancestry performance of PUMAS-ensemble PRS for blood lipid traits in UKB.** We trained PUMAS-EN and PUMAS-SL PRS models for AFR, EAS, EUR, and SAS ancestries for traits **(A)** LDL cholesterol **(B)** logTG (**C)** TC.

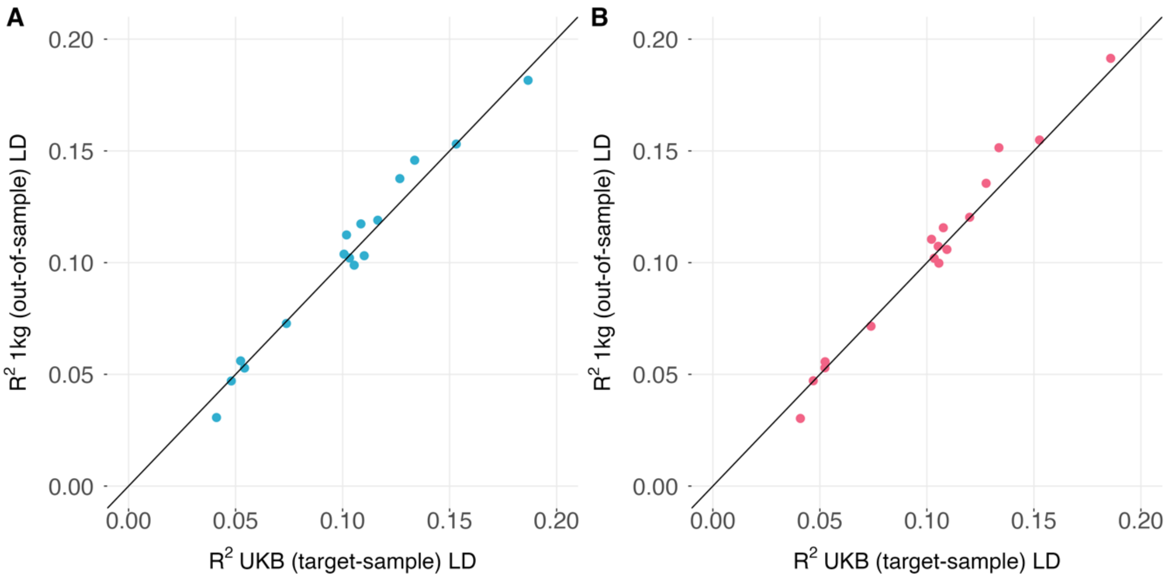


**Fig S5**. **Comparing PUMAS-ensemble models trained using target-sample LD (UKB) and external LD (1KG) in GLGC lipid PRS analysis**. Both LD panels match ancestries of the target dataset. Results for PUMAS-EN and PUMAS-SL are shown in panels A and B, respectively. Y-axis: $R^2$ from PUMAS-ensemble trained using 1KG LD; X-axis: $R^2$ from PUMAS-ensemble trained using UKB LD. The diagonal line is shown in each panel.

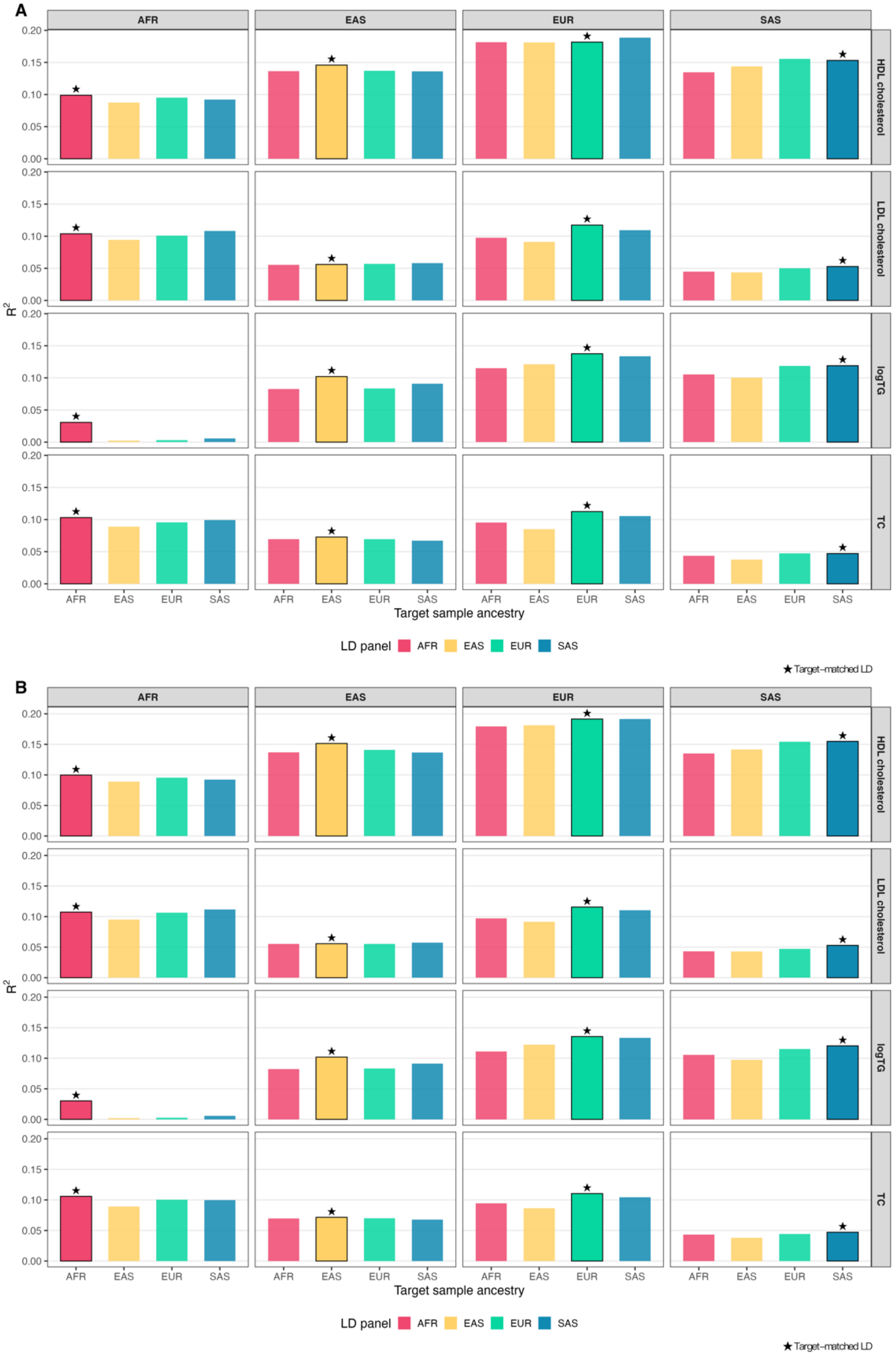


**Fig S6**. **Comparing PUMAS-ensemble models trained using external LD (1KG) with and without matching ancestry of the target dataset in GLGC lipid PRS analysis**. Performance of PUMAS-EN and PUMAS-SL for 4 lipid traits (row) and 4 target datasets of different ancestries (column) are shown in panels A and B, respectively. LD panels of AFR (red), EAS (yellow), EUR (cyan), and SAS (blue) ancestries were used to train PUMAS-ensemble models. Y-axis (within each cell): PUMAS-ensemble $R^2$; X-axis (within each cell): LD panel used to train PUMAS-ensemble. Performance of PUMAS-ensemble trained using ancestry-matched external LD (1KG) is highlighted with asterisk.
